# *Vibrio parahaemolyticus* metabolically adapts while tempering the host immune response during host cell invasion

**DOI:** 10.64898/2026.08.19.745812

**Authors:** Yayun Zheng, Chenglong Sun, Sukhithasri Vijayrajratnam, Jananee Jaishankar, Lisa N. Kinch, Zhijian J. Chen, Kim Orth

## Abstract

*Vibrio parahaemolyticus* (*V. para*) is an enteric pathogen that establishes a protected intracellular niche using its second type III secretion system. However, how this bacterium adapts to the host cytoplasm while overcoming cellular defenses has remained unclear. To define these mechanisms, we performed dual-transcriptomic profiling of both pathogen and host during invasion, intracellular replication, and late infection. Our analyses revealed extensive metabolic reprogramming by *V. para*, including induction of diverse nutrient transporters and metabolic pathways. We discovered that because mammalian cells are auxotrophic for aromatic amino acids, *V. para* must activate its own unique biosynthetic machinery, a requirement that proved essential for intracellular growth. Infected host cells mounted a sustained NF-κB response. Both heightened NF-κB activation and disruption of canonical NF-κB signaling restricted bacterial expansion, indicating that *V. para* exploits a finely tuned “*Goldilocks*” level of immune signaling to promote survival and replication. Together, these findings uncover fundamental metabolic and immune adaptations that drive pathogenesis.

## Introduction

*Vibrio parahaemolyticus* (*V. para*), an enteric pathogen found in marine and estuarine environments, represents a leading cause of seafood-borne gastroenteritis in humans worldwide^1^. With rising sea surface temperatures, *V. para* can survive in expanded geographic areas where they were previously rare, increasing the risk for human consumption^2,3^. Rising temperatures have also been associated with increased pathogenicity of environmental *V. para* isolates as well as their enhanced resistance to environmental stressors^4,5^. These factors have contributed to *V. para* being designated as an emerging foodborne pathogen^6,7^ and highlights the need for understanding its virulence mechanisms associated with human infection.

*V. para* encodes two type III secretion systems (T3SS): the first, T3SS1, is found in all strains; and the second, T3SS2, is associated with clinical isolates and disease. T3SSs encode a needle-like apparatus used to inject a set of effectors into host epithelial cells to manipulate host signaling for the benefit of the pathogen. For invasion, the T3SS2-encoded VopC deamidase effector activates the small GTPase Rac to induce membrane ruffling of host epithelial cells and promote bacterial uptake^8–10^. After invasion, *V. para,* escapes into the cytosol, where it replicates to approximately 200–300 bacteria per host cell within 7-8 hrs^11^. During this intracellular replication phase, *V. para* uses additional virulence factors to manipulate host-cell responses and promote intracellular survival, including a constitutively secreted lipase that weakens the host cell plasma membrane to aid *V. para* egress^12^. Additionally, *V. para* modifies its lipid A to evade recognition by the host innate immune systems^13^. Despite this mechanistic knowledge of *V. para* interactions with its human host, how *V. para* coordinates these strategies to remodel the intracellular host environment and support its survival and replication remains poorly understood.

Epithelial cells form the first barrier against enteric pathogens and provide a physiologically relevant context for understanding this enteric bacterial pathogenesis and host barrier defense^14^. As a representative example of host cell invasion, we used Caco-2 cells, a human colorectal adenocarcinoma immortalized cell line, for infections with *V. para*. Approximately 4-10% of Caco-2 cells are invaded with *V. para*, followed by replication and escape of 300 bacteria over about 7 hours^15,16^. To gain transcriptomic insight over the course of an infection cycle, we chose to use dual RNA-seq, as this method has emerged as a powerful approach for studying infection from both microbial and host perspectives^17^. Its landmark application to *Salmonella* infection showed that simultaneous profiling can reveal infection-induced bacterial regulatory programs alongside the accompanying host transcriptional response^18^. Subsequent work extending dual RNA-seq to other intracellular pathogens in infected cells, tissues, and whole-organism models further demonstrated its utility to uncover physiologically informative, infection-specific transcriptional states for both the host and pathogen^19–23^. In the case of *V. para*, an enteric bacterium with a transient but productive intracellular phase in non-phagocytic epithelial cells, such an approach provides a time-resolved framework to connect bacterial adaptation, host responses, and progression through the complete intracellular infection cycle.

To define this process, we applied time-resolved dual transcriptomic analysis to Caco-2 epithelial cells invaded with a modified clinical isolate of *V. parahaemolyticus* RIMD2210633 (CAB2; deleted for hemolysins and T3SS1 expression) at defined infection stages^16^. Our results reveal that intracellular *V. para* activates layered transcriptional programs across the infection cycle including early adjustment of nutrient-acquisition systems and metabolic pathways. During late infection, *V. para* induces surface-associated programs, including flagellar machinery, consistent with a transition from intracellular expansion toward egress or dissemination. Importantly we observe invasion induces modest transcriptional changes in the host that includes a moderate induction of the protective components in NF-κB pathway during intracellular *V. para* infection. In summary, our study reconstructs the host–pathogen transcriptional trajectory of a complete intracellular infection cycle and reveals how *V. para* couples metabolic adaptation with host immune-state modulation to establish a replication-permissive epithelial niche.

## Results and Discussion

### Regulated transcriptional changes observed for host and pathogen during intracellular infection

To investigate the physiological response of invading bacteria and host cells, we infected the human intestinal epithelial cell line Caco-2 with a GFP-expressing *V. para* CAB2 strain pretreated with the bile acid taurodeoxycholate (TDC) to induce T3SS2-mediated internalization^8^. We selected three time points after infection (1.5 h, 4 h and 6.5 h) to capture the progression of *V. para* from early entry to rapid intracellular replication and pre-egress/egress, respectively. Intracellular bacterial replication at these time points was monitored by CFU measurements (**Fig. 1a**), showing low bacterial numbers during early infection, a marked increase during rapid intracellular replication, and a subsequent decline at the late pre-egress/egress stage. This intracellular infection time course was established in our previous studies^8,16,24^, and here we provide corresponding microscopy images illustrating these stages in Caco-2 cells (**Fig. 1a**).

**Figure 1.**
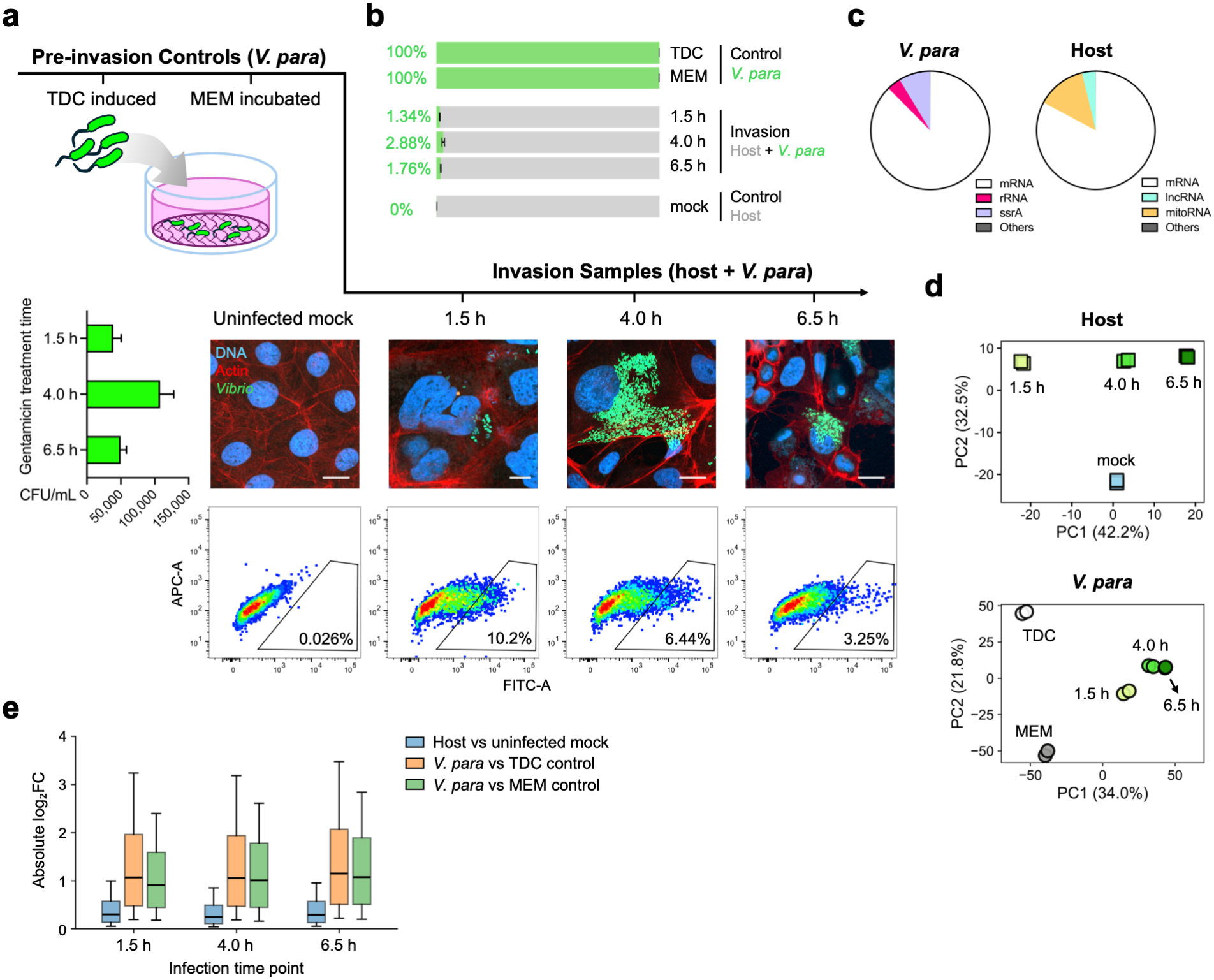
Dual RNA-seq analyses reveal transcriptomic profiles of both host and bacteria during *V. para* infection. **(a)** Dual RNA-seq workflow. Caco-2 cells invaded by GFP-expressing *V. para* were enriched by FACS. Bacterial samples were collected under two pre-invasion conditions (TDC and MEM), and infected cells were collected at three time points following gentamicin treatment to eliminate extracellular bacteria. Lower panels show intracellular CAB2 loads at the RNA-seq sampling time points (CFU; mean ± s.d., *n* = 3; left), representative microscopy images (scale bar, 10 μm; right), and FACS plots showing GFP-positive gating and percentages. **(b)** Transcript composition of infected-cell and control libraries (mean ± s.d., *n* = 2). TDC, *V. para* grown in MLB after TDC induction; MEM, induced bacteria incubated in cell culture medium; mock, uninfected Caco-2 cells. **(c)** Representative mapping statistics for bacterial (left) and host (right) reads from the 4 h infected-cell library. Fractions below 1% are grouped as “Others”. **(d)** Principal component analysis (PCA) of host (upper) and bacterial (lower) transcriptomes (two biological replicates per condition). Host PCA used variance-stabilized expression data excluding mitochondrial genes; bacterial PCA used TPM values excluding rRNAs and *ssrA*. Axis labels indicate the variance explained by each PC. **(e)** Distributions of absolute log2(fold change) relative to the indicated controls for genes with DESeq2 baseMean ≥ 10 across all comparisons within each species. The same genes were used across time points. Boxes indicate interquartile ranges, center lines indicate medians, and whiskers extend from the 10^th^ to 90^th^ percentiles.

To obtain invaded cells for transcriptomic profiling, *V. para*-infected cells were identified by intracellular GFP signal and collected by fluorescence-activated cell sorting (FACS)^18^ (**Fig. 1a**). To distinguish infection-associated transcriptional changes from baseline host and bacterial states, we included an uninfected mock control for the host cells, and two pre-invasion bacterial reference conditions: 1) *V. para* after 90 min of T3SS2 induction with the bile acid TDC^25,26^ and 2) TDC-induced *V. para* further incubated in MEM to mimic exposure to the host-cell culture medium (**Fig. 1a**).

Because the number of infected cells sorted by FACS was limited (around 100,000 cells per sample), we adopted a low-input library construction workflow based on a tagmentation-based template-switch protocol^27^. This strategy was combined with a customized rRNA depletion pipeline to remove both host and bacterial rRNAs through hybridization-based digestion^28^ (see **Methods**; **Fig. S1**). We then assessed whether the resulting libraries captured both host and bacterial transcripts with sufficient coverage. Across the three infection time points, bacterial transcripts accounted for only 1–3% of total transcripts (**Fig. 1b**). This low fraction nevertheless provided near-complete coverage of the intracellular *V. para* transcriptome, with transcripts detected from 4,710 of 4,991 annotated genes across the time course. Moreover, more than 80% of mapped reads in both host and bacterial libraries mapped to mRNA, indicating efficient rRNA depletion and yielding abundant informative transcripts for downstream analysis (**Fig. 1c**).

We next examined whether the dataset reflected broad transcriptional differences between intracellular infection samples and their reference conditions. Principal component analysis of the host transcriptome showed that mock controls separated from infected samples along the second principal component (PC2, 32.5%), while infected host-cell samples were ordered by infection time (1.5 h, 4 h and 6.5 h) along the first principal component (PC1, 42.2%), indicating progressive remodeling of the host transcriptome during infection (**Fig. 1d**). In the bacterial dataset, intracellular *V. para* samples were clearly separated from both pre-invasion references (TDC and MEM) along both PC1 (34.0%) and PC2 (21.8%). Differences among the intracellular time points were more modest, with the 1.5 h samples showing a shift relative to the 4 and 6.5 h samples, whereas the pre-invasion reference samples occupied distinct positions from the intracellular samples along both PC axes (**Fig. 1d**). Together, these analyses show that infection status represents a major source of transcriptomic variation in both host cells and bacteria, while also revealing progressive transcriptional changes across the course of intracellular infection.

Overall, our dual RNA-seq pipeline successfully profiled changes in both bacterial and host transcriptomes throughout the intracellular infection cycle. Interestingly, relative to their respective control or reference conditions, gene-wise expression changes in host cells were markedly less pronounced than those observed in *V. para* across all post-invasion time points: in host cells, approximately 7–10% of genes changed by more than two-fold relative to the mock control across the three infection time points; by contrast, this proportion reached 46–55% in *V. para* relative to the pre-invasion TDC or MEM references (**Fig. 1e**). Although comparisons of the transcriptional responses between the two species should be interpreted with some caution given differences in baseline conditions and the nature of gene regulation in prokaryotes and eukaryotes, the markedly smaller magnitude of global changes in host cells nevertheless indicates a relatively limited transcriptional response to intracellular infection.

Using this dataset, we next investigated bacterial adaptation and host responses during the intracellular invasion time course.

### *V. para* activates distinct temporal profiles for gene clusters during infection

We first defined the temporal patterns of bacterial gene expression during intracellular infection through the three infection time points by calculating the expression trajectory of each gene compared to the pre-invasion MEM sample. Genes were grouped by their expression trajectories using unsupervised k-means clustering, and the resulting clusters were visualized by Uniform Manifold Approximation and Projection (UMAP) (**Fig. 2a**). Six clusters exhibited characteristic temporal expression patterns representing three major behaviors: repression after invasion (**Fig. 2b**, repression), transient activation during specific stages of infection at 1.5 h, 4 h and 6.5 h (**Fig. 2b**, unimodal activation), and sustained activation throughout intracellular growth after 1.5 h or 4 h (**Fig. 2b**, plateau activation). Cluster 7 lacked a clear shared temporal expression pattern and was therefore categorized as “unclassified” (**Fig. 2b**).

**Figure 2.**
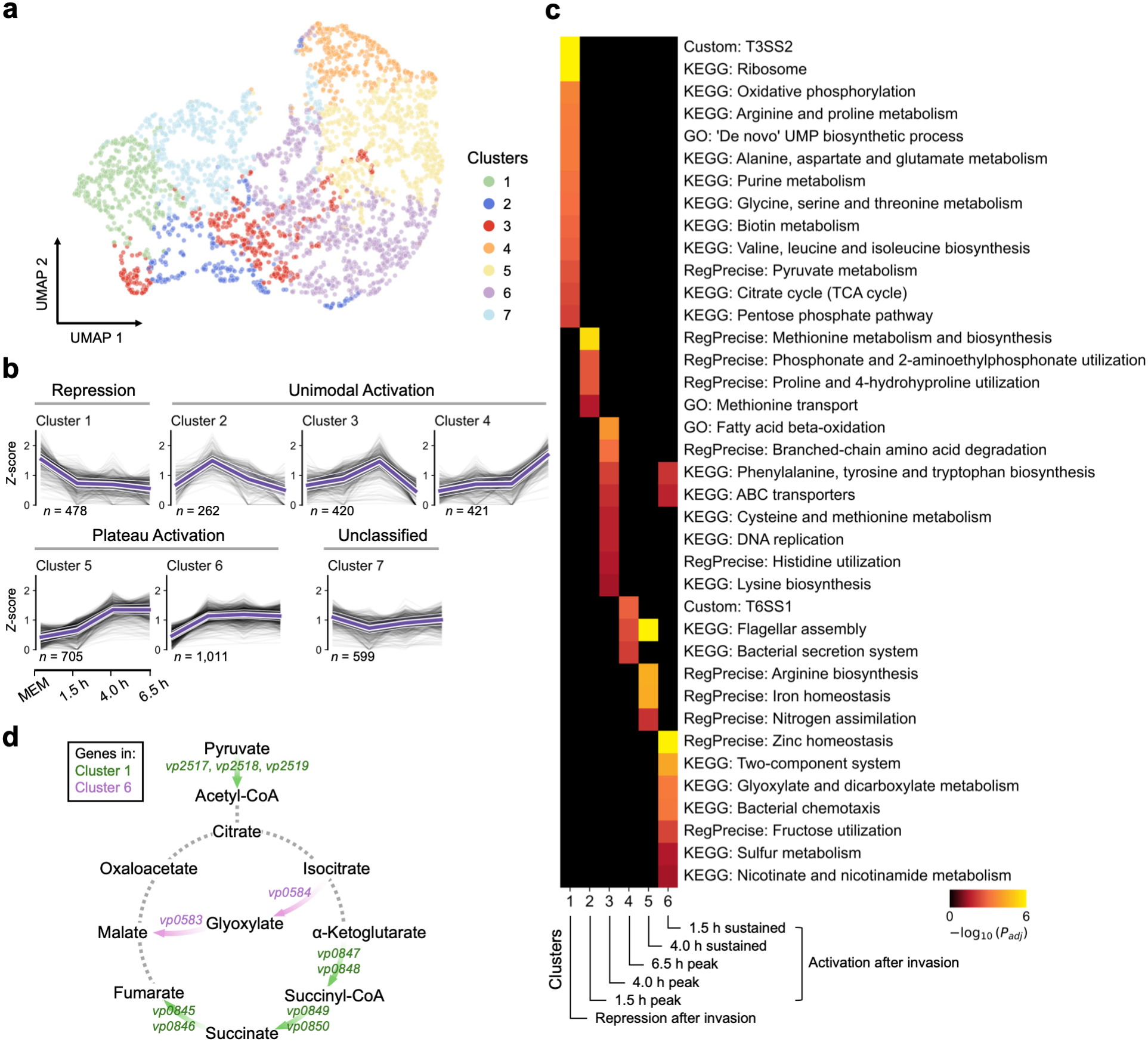
*V. para* genes exhibit dynamic expression patterns during intracellular infection. **(a)** UMAP visualization of all detected *V. para* genes. Each dot represents one gene and is colored according to its assigned expression cluster as illustrated in **b**. **(b)** *Z*-score-normalized expression profiles of genes within each cluster across the MEM pre-invasion condition and intracellular infection time points (1.5, 4.0, and 6.5 h). Faint gray lines indicate individual gene trajectories, and the bold purple line represents the cluster mean. Purple shading denotes the interquartile range, with white boundary lines marking the 25^th^ and 75^th^ percentiles. The number of genes in each cluster is indicated below each plot. **(c)** Heatmap showing representative enriched functional terms for each cluster in b, identified by hypergeometric tests. Functional annotations include KEGG pathways (KEGG), GO Biological Process (GO), RegPrecise-predicted regulons (RegPrecise), and manually curated gene sets (Custom), with their sources indicated by prefixes before each term. Heatmap colors indicate Benjamini–Hochberg-adjusted *P* values for GO terms and nominal *P* values for the remaining functional gene sets (cutoff < 0.05). Redundant, highly overlapping, or parent/child terms representing largely the same gene sets are omitted for clarity. Cluster identities are indicated below each column. **(d)** Simplified schematic of the TCA cycle and glyoxylate shunt. Arrows and gene labels are colored according to the legend and follow the same color scheme as in **a**. Other reactions are shown as gray dashed lines.

To identify the biological functions associated with these bacterial transcriptional programs, we performed functional enrichment analysis for each expression trajectory cluster (**Fig. 2c; Table S1**). Given the limited functional annotation coverage for *V. para*, we incorporated KEGG pathways^29^, GO Biological Process annotations^30^, RegPrecise-predicted regulons^31^, and manually curated gene sets covering bacterial secretion systems and pathogenicity islands. Together, these complementary annotations provided broader functional coverage, facilitating a more comprehensive interpretation of the biological functions represented by each cluster (see **Methods**).

Genes repressed after invasion (cluster 1) were enriched for T3SS2, protein translation (ribosome) and pathways involved in central carbon metabolism (TCA cycle, pentose phosphate pathway, and pyruvate metabolism), energy production (oxidative phosphorylation), and amino acid and nucleotide metabolism (**Fig. 2c**). This broad transcriptional repression suggests that *V. para* downregulates growth-associated metabolic and biosynthetic programs, along with invasion-associated secretion machinery, as it adapts to the intracellular environment. In contrast, genes activated throughout infection (cluster 6) were enriched for environmental sensing (two-component systems and bacterial chemotaxis) and nutrient acquisition (ABC transporters). Additional enrichment was observed for fructose utilization, sulfur metabolism, aromatic amino acid biosynthesis, and zinc homeostasis, suggesting sustained requirements for these functions during intracellular bacterial growth (**Fig. 2c**). Notably, cluster 6 was also enriched for glyoxylate and dicarboxylate metabolism (**Fig. 2c**), including the key glyoxylate shunt genes *aceA* and *aceB*. Together with the repression of TCA cycle genes in cluster 1, this pattern suggests a possible shift toward the glyoxylate shunt during intracellular growth (**Fig. 2d**).

Several pathways showed stage-specific activation (**Fig. 2c**). At 1.5 h post-infection, methionine transport and metabolism were selectively enriched, suggesting an early adaptation response immediately after host-cell entry. At 4 h post-infection, during the rapid intracellular replication phase, *V. para* showed strong enrichment of genes involved in fatty acid β-oxidation, metabolism and biosynthesis, and ABC transporters, indicating increased nutrient acquisition and metabolic remodeling. DNA replication genes were also enriched at this time point, further supporting active bacterial proliferation. At 6.5 h post-infection, flagellar assembly and bacterial secretion systems, including T6SS1, became enriched, suggesting a late transition from intracellular propagation toward motility- and secretion-associated processes that may facilitate bacterial dissemination from host cells.

Together, these results indicate that intracellular adaptation of *V. para* is organized into multilayered transcriptional programs. These include: 1) post-invasion repression of translation, biosynthesis, and central metabolism, consistent with reduced nutrient availability within host cells; 2) sustained activation of metabolite-sensing and nutrient-acquisition systems; and 3) stage-specific shifts in bacterial programs, from early adaptation to mid-stage replication and, ultimately, to late induction of motility and secretion pathways in preparation for egress. Moving forward, we focused on two major infection-associated processes revealed by this temporal analysis: nutrient acquisition and metabolic remodeling during intracellular growth, and activation of secretion- and motility-associated pathways during late infection.

### Intracellular *V. para* shifts arginine metabolism from catabolism toward concurrent uptake and biosynthesis

Intracellular replication was accompanied by extensive metabolic remodeling, particularly of amino acid metabolism. Among these changes, the arginine and proline metabolism pathway was enriched among genes repressed after host-cell entry (**Fig. 2c**). Examination of the underlying genes revealed that several genes preferentially expressed before invasion were involved in arginine catabolism, including *astA* and *astD* of the arginine succinyltransferase pathway, together with genes involved in arginine-dependent polyamine biosynthesis (**Table S1**). These observations indicate a shift away from arginine catabolism and polyamine production following host-cell entry, suggesting a broader reorganization of arginine utilization during intracellular growth.

To further resolve this transition, we examined the predicted ArgR regulon, which includes operons involved in arginine catabolism, biosynthesis, and uptake. ArgR senses L-arginine and binds conserved ARG-box sequences in the promoters of its target operons, repressing genes involved in arginine biosynthesis while promoting the expression of specific arginine catabolic operons in some bacteria^32,33^. A comparative genomic reconstruction identified a predicted ArgR regulon in *V. para*^31^, whose genes separated into two opposing temporal expression programs (**Fig. 3a**). Arginine catabolic genes were preferentially expressed before invasion and repressed after host-cell entry, whereas biosynthetic genes were induced during rapid intracellular replication at 4 h and remained elevated at 6.5 h. This reciprocal pattern indicates a shift from the use of arginine as a catabolic substrate toward increased arginine biosynthesis, consistent with greater anabolic demand during rapid intracellular replication.

**Figure 3.**
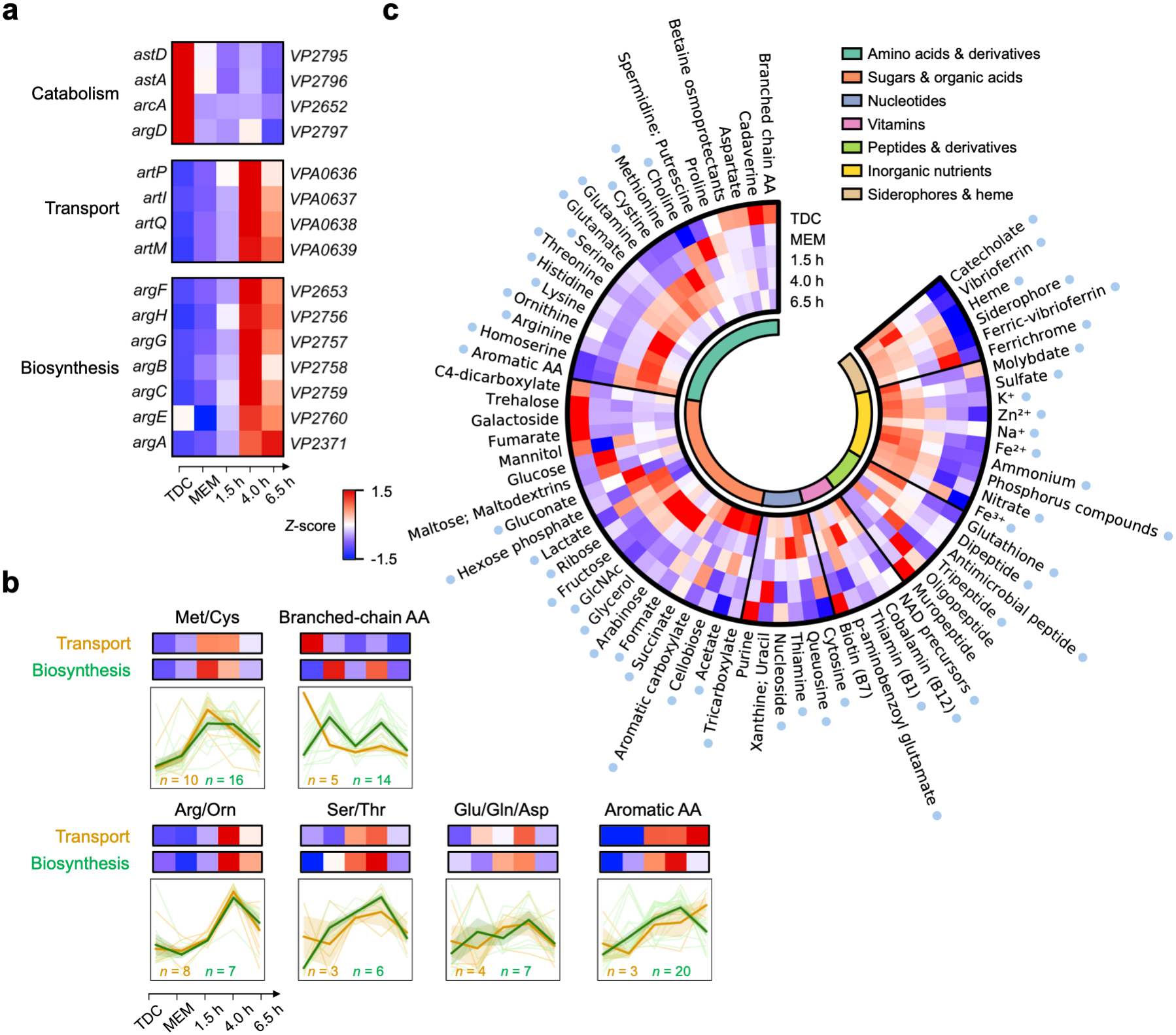
Coordinated remodeling of amino acid metabolism and nutrient acquisition during intracellular *V. para* infection. **(a)** *Z*-score-normalized expression profiles of ArgR-regulated genes across pre-infection and intracellular infection conditions. Rows represent individual genes, with gene names shown on the left and locus tags on the right. Genes are arranged into three functional sections from top to bottom: arginine catabolism, transport, and biosynthesis. **(b)** Normalized expression patterns of amino acid transporter and biosynthetic genes across all conditions, grouped into six amino acid subgroups. Heatmaps show averaged *Z*-score-normalized expression levels for genes within each subgroup and functional category. Line plots show individual gene trajectories as faint lines, with the thick colored line representing the group average. The number of genes included in each group is indicated. **(c)** Heatmap showing expression patterns of transporters categorized by their transported substrates across pre-infection and infection conditions. Each column represents one substrate and shows the average *Z*-score-normalized expression of transporters assigned to that substrate. Rows indicate different conditions. The inner annotation circle indicates the broader substrate category. Blue dots mark substrates with intracellular activation, defined as either sustained elevation across two consecutive intracellular time points or a single peak across the three intracellular time points.

The predicted ArgR regulon in *V. para* also includes the *artPIQM* genes, which encode an arginine ABC transporter^31^ (**Fig. 3a**). Their temporal expression closely paralleled that of the biosynthetic genes, with induction after host-cell entry, peak expression at 4 h, and sustained elevation at 6.5 h, consistent with their assignment to the corresponding temporal cluster in **Fig. 2c**. The coordinated induction of arginine biosynthesis and uptake suggests that intracellular *V. para* uses complementary strategies to maintain arginine availability during rapid replication.

Beyond arginine, coordinated temporal regulation of transport and biosynthesis was also observed for other amino acids. Another example was provided by methionine, whose transport and biosynthetic genes were both enriched in the early-response cluster that peaked at 1.5 h (**Fig. 2c**). To determine whether such coordination extends more broadly across amino acid metabolism, we examined representative groups including sulfur-containing, branched-chain, arginine/ornithine-related, hydroxyl-containing, central nitrogen metabolism-related, and aromatic amino acid (**Fig. 3b**). Except for branched-chain amino acids, transporter and biosynthetic genes within each group showed similar temporal profiles, characterized by induction after host-cell entry. Sulfur-containing amino acid transport differed in timing, peaking earlier at 1.5 h, consistent with the cluster-based enrichment result (**Fig. 2c**), whereas the other transporter and biosynthetic programs generally reached peak expression during rapid replication at 4 h (**Fig. 3b**). Together, these findings highlight a shift in amino acid metabolism from catabolism during pre-invasion growth toward coordinated biosynthesis and nutrient uptake during intracellular replication, consistent with adaptation to a more nutrient-limited intracellular environment.

### Remodeling of nutrient acquisition during intracellular replication of *V. para*

The coordinated induction of amino acid transport systems prompted us to examine how nutrient acquisition is remodeled more broadly during intracellular growth. To define this program more systematically, we generated a curated profile of nutrient-uptake systems, including predicted transporters and ABC transporter complexes (**Table S2**). Because existing annotations are incomplete and often lack substrate-level resolution, we used a substrate-oriented list of *E. coli* transporters^34^ as a reference to identify potential corresponding homologs by BLAST. These annotations were then manually refined using UniProt^35^ and RefSeq annotations^36^ (see **Methods**). This framework allowed us to classify nutrient-uptake systems by predicted substrate class and examine their expression across the infection time course.

Using this substrate-based transporter profile, we compared intracellular bacteria with two pre-invasion bacteria references (**Fig. 1a**, TDC bacterial culture medium and MEM host-cell medium). Relative to these reference conditions, transporters predicted to be associated with amino acids, carbon substrates, nucleoside- and cofactor-related metabolites, inorganic nutrients, metal ions, and iron acquisition were induced during intracellular infection. This pattern suggests that intracellular *V. para* increases its capacity to acquire multiple nutrient classes when transitioning from *in vitro* culture settings to physiological context of the host cell (**Fig. 3c**, highlighted dots).

Within this broader response, nutrient-acquisition programs showed a staged pattern of activation (**Fig. 3c**). Most amino acid transport systems peaked at 4 h, whereas those associated with sulfur-containing amino acids showed the strongest early response, peaking at 1.5 h, consistent with both the cluster-based analysis (**Figs. 2b, c**) and the pathway-level analysis (**Fig. 3b**). Carbon-source transporters showed a similarly staged pattern: glucose-associated transporters peaked during pre-invasion incubation, whereas intracellularly induced systems from 1.5 h to 4 h were linked to gluconate, hexose phosphate, fructose/PTS-associated sugars, N-acetylglucosamine, lactate, and ribose. Several of these substrates have been implicated in the intracellular metabolism^37–40^ or host-associated nutrient acquisition^41,42^, suggesting that intracellular *V. para* may access host-available carbon sources to support growth. In comparison, inorganic nutrient transporters showed stronger expression at 4 h or 6.5 h, suggesting that these resources may be increasingly required in the later phase of the intracellular life cycle (**Fig. 3c**). Siderophore-associated iron-acquisition systems followed a distinct pattern, with elevated expression beginning during pre-invasion incubation and persisting throughout intracellular infection (**Fig. 3c**; **Fig. S2**). The potential metabolic implications of these sulfur- and iron-acquisition responses are discussed further below.

### Induction of aromatic amino acid biosynthetic pathways essential for *V. para* proliferation

During early and mid-stage intracellular infection, *V. para* showed increased expression of genes involved in phenylalanine, tyrosine, and tryptophan biosynthesis (**Fig. 2c**; **Fig. 3b**). This aromatic amino acid biosynthetic pathway was of particular interest because bacteria can synthesize these amino acids *de novo*, while mammals lack enzymes in the shikimate pathway and are auxotrophic for their acquisition. In *V. para*, genes encoding the upstream steps of the shikimate pathway showed heterogeneous expression patterns, with several genes induced more strongly during pre-invasion incubation in MEM than during intracellular growth. By contrast, genes in the chorismate-utilizing branches, particularly those involved in aromatic amino acid biosynthesis, tended to show higher expression during intracellular growth (**Fig. 4a**). These observations suggest preferential activation of downstream aromatic amino acid biosynthesis during intracellular replication.

**Figure 4.**
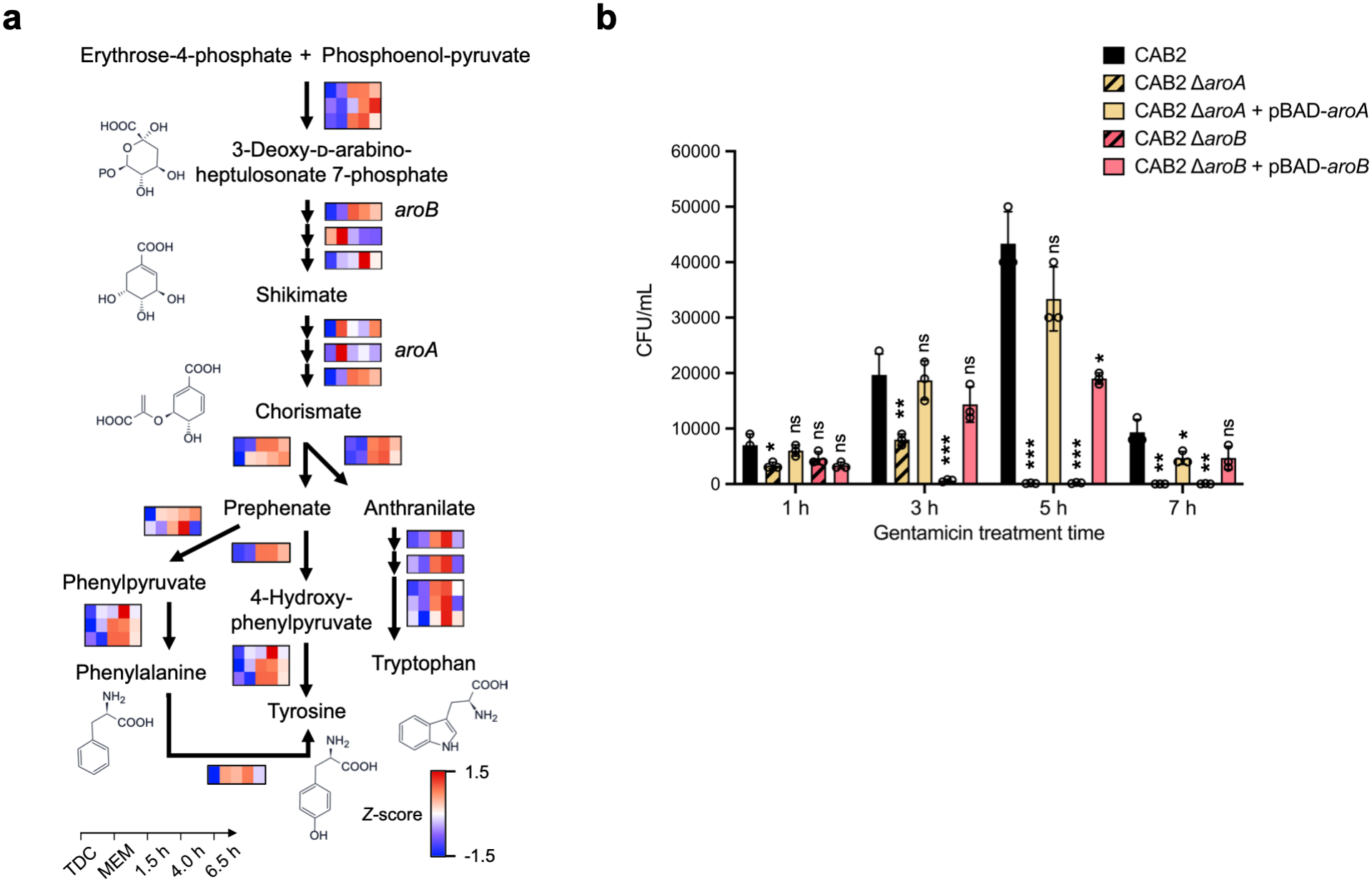
Intracellular *V. para* depends on aromatic amino acid biosynthesis for replication. **(a)** Pathway schematic showing expression dynamics of genes involved in chorismate-derived aromatic amino acid biosynthesis. Arrows indicate individual enzymatic reaction steps, and adjacent heatmap rows show the normalized expression profiles of corresponding genes across five conditions. Multiple rows assigned to the same step indicate redundant predicted to catalyze the same reaction. **(b)** Intracellular replication of *V. para* CAB2, Δ*aroA*, Δ*aroB*, and their corresponding complemented strains in Caco-2 cells. Cells were infected for 90 min and then incubated in gentamicin-containing medium. Intracellular bacterial loads were determined at the indicated time points post-gentamicin treatment. Note that throughout this study, we used our standard time course (1, 3, 5, and 7 h) to assess intracellular replication dynamics, rather than the streamlined time points (1.5, 4, and 6.5 h) selected for dual RNA-seq. Data are shown as mean ± s.d. (*n* = 3 biological replicates). Statistical significance was assessed by Student’s *t*-test relative to CAB2 at each time point and is indicated above each column: ns, *P* ≥ 0.05; *, *P* < 0.05; **, *P* < 0.01; ***, *P* < 0.001.

To determine whether chorismate biosynthesis is functionally required for intracellular growth, we independently deleted two shikimate pathway genes (*aroA* or *aroB*) that are required for chorismate production. Loss of either gene strongly impaired intracellular replication while restoration of intracellular replication was observed with the mutant strain complimented with the corresponding deleted gene (**Fig. 4b**; **Fig. S3**). These results validate our hypothesis that *de novo* production of chorismate-derived aromatic metabolites is required for efficient intracellular replication of *V. para*.

### *V. para* secretion systems display dynamic patterns across phases of intracellular infection

Having characterized metabolic and nutrient-acquisition programs that support intracellular replication, we next examined the late-stage secretion and motility pathways identified by the temporal analysis (**Fig. 2c**), with a focus on individual secretion systems and flagellar components. Bacterial secretion systems showed distinct temporal patterns rather than uniform induction across the course of intracellular infection (**Fig. 5a**; **Table S3**). Consistent with the enrichment of T3SS2 among genes repressed after invasion (cluster 1; **Fig. 2c**), T3SS2 genes responsible for cell invasion were expressed most highly in the TDC-induced pre-invasion state. After host-cell entry, expression of T3SS2 structural genes, chaperones and effectors progressively decreased. This decrease suggests that sustained T3SS2 expression depends on continued exposure to the bile salt TDC, the T3SS2 inducing ligand^43^. T3SS2 is likely dispensable after bacterial internalization, especially during intracellular replication. T3SS1 genes showed minimal expression consistent with the T3SS1-disabled genetic background of this strain^8^, providing an internal validation of our dual RNA-seq analysis. Intriguingly, T2SS genes remained expressed during intracellular infection (**Fig. 5a**). The T2SS secretes extracellular substrates that promote nutrient acquisition, biofilm formation and pathogenicity, and is increasingly recognized as a driver of virulence^44^. Additionally, VPA0226, a T2SS-associated lipase required for *V. para* escape from host cells^12^, had increased gene expression after host-cell entry, matching the timing expected for a late-stage egress factor. Consistent with reports that T6SS1 is responsive to TDC^45^, T6SS1 genes were highly expressed in the TDC-induced pre-invasion state and decreased after host-cell entry, showing a pattern partially related to T3SS2 (**Fig. 5a**). As T6SS1 is best known as an antibacterial weapon that delivers toxic effectors into neighboring bacterial competitors, its downregulation after host-cell entry is not unexpected^46^. However, several T6SS1 genes showed a modest increase at the 6.5 h, consistent with T6SS1 enrichment in cluster 4 (**Fig. 2c**), while T6SS1 effector genes remained relatively stable across intracellular infection. These patterns suggest that T6SS1 may remain poised after invasion. Together, these patterns indicate that secretion programs are temporally partitioned during intracellular infection: T3SS2 represents a pre-invasion-induced program, T2SS expression is maintained with increased expression of the egress-associated effector VPA0226, and T6SS1 may continue to respond to infection-associated cues after invasion.

**Figure 5.**
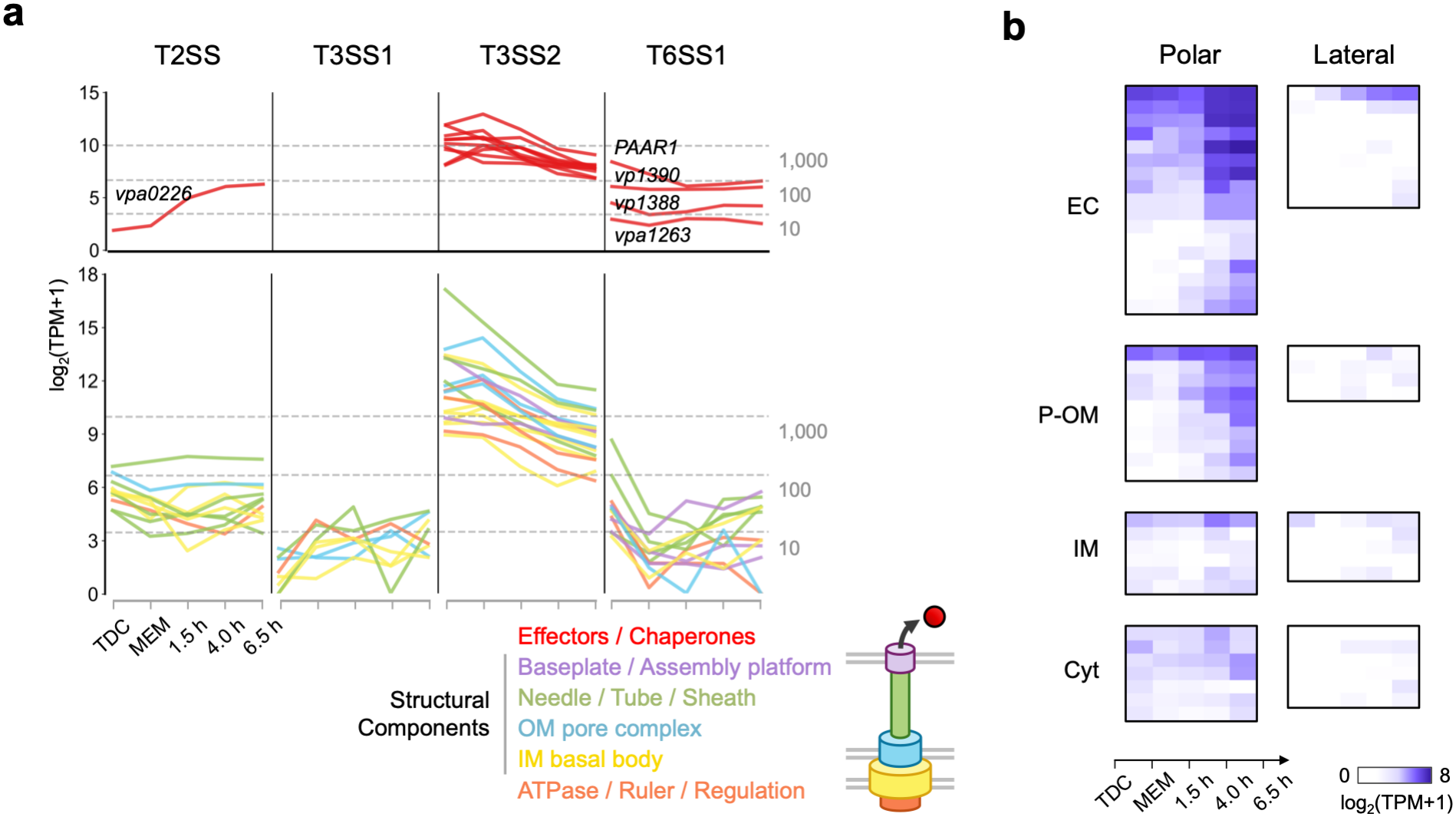
Intracellular infection induces stage-specific expression of *V. para* secretion and flagellar systems. **(a)** Expression profiles of genes from different secretion systems across pre-infection and intracellular infection conditions. For each secretion system, effectors and chaperones are shown in red in the upper panel, whereas structural and other system-associated components are shown in the lower panel and color-coded according to the functional categories indicated in the schematic. Effector names are labeled in the plots except for T3SS2; complete annotations are provided in **Table S3**. Each line represents the mean expression profile of one gene across the indicated conditions (*n* = 2 biological replicates). **(b)** Heatmaps showing expression profiles of polar and lateral flagellar genes across pre-infection and intracellular infection conditions. Each row represents one gene, and each column shows the mean expression level at the indicated conditions or infection time points (*n* = 2 biological replicates). Genes are grouped according to their predicted subcellular localization. EC, extracellular; P-OM, periplasmic/outer membrane; IM, inner membrane; Cyt, cytoplasmic.

Flagellar genes showed a pronounced late-stage pattern (**Fig. 5b**). For the polar flagellum, expression changes were not uniform across the entire flagellar apparatus. Instead, induction was biased toward genes encoding extracellular or surface-exposed components: genes encoding distal flagellar structures (flagellins, the cap and hook-associated components) progressively increased during intracellular infection, whereas genes encoding inner-membrane, export or cytoplasmic components remained comparatively stable. Genes encoding periplasmic or outer-membrane components showed intermediate induction. Lateral flagellar genes showed a similar trend, although both their induction and absolute expression levels were lower than those of the polar flagellar system. Together, these patterns suggest selective remodeling of the flagellar program during late intracellular infection, with preferential induction of surface-associated components.

### NF-κB signaling exhibits a *Goldilocks Principle* for intracellular survival of *V. para*

Having defined the major bacterial programs activated during intracellular infection, we next asked how host cells respond to intracellular *V. para*. To identify coordinated pathway-level host responses, we performed functional gene set enrichment analysis (GSEA) by comparing infected host cells with uninfected mock controls at each time point across the infection course. We used the Hallmark gene set collection to evaluate host function, which consolidates related biological processes into 50 coherent gene sets with reduced redundancy and heterogeneity^47^. Among the top enriched pathways, the strongest positive enrichment was observed for inflammatory programs during early and mid-stage infection, including interferon-γ response, TNF-α signaling via NF-κB, and inflammatory response (**Fig. 6a, b**). These results suggest that intracellular *V. para* infection induces a host inflammatory transcriptional program, with NF-κB signaling emerging as a prominent response. In addition to these programs, an apoptosis hallmark was enriched specifically at 1.5 h (**Fig. 6a**), and further examination of apoptosis-related genes revealed increased expression of negative regulators of apoptotic signaling (**Fig. 6c**). Together, these results indicate a host response characterized by inflammatory signaling and concurrent induction of anti-apoptotic genes.

**Figure 6.**
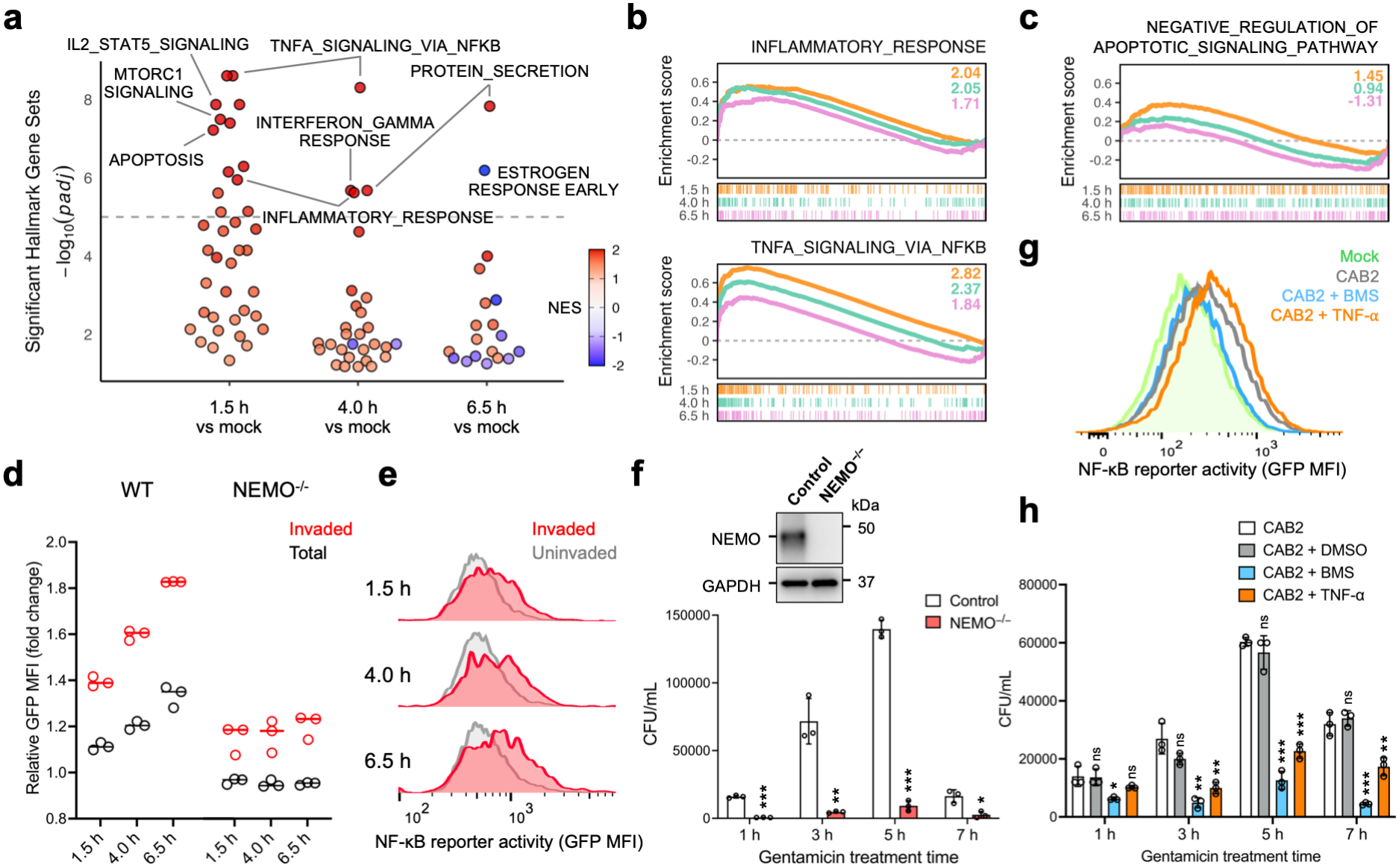
*V. para* infection activates host NF-κB signaling during intracellular replication. **(a)** Significant hallmark gene sets from GSEA of infected versus mock-infected cells across time points (nominal *P* < 0.05; FDR *q* < 0.25). Dots represent gene sets; position indicates time and −log10(adjusted *P*), and color indicates normalized enrichment score (NES). The dashed line marks −log10(adjusted *P*) = 5; selected pathways above this threshold are labeled. **(b)** GSEA running-score plots of NF-κB-related pathways. NES values are indicated; orange, 1.5 h; teal, 4 h; pink, 6.5 h. **(c)** GSEA plots of GOBP pathways involved in negative regulation of apoptotic signaling. Colors as in **b**. **(d)** NF-κB reporter activity (GFP mean fluorescence intensity, MFI) normalized to mock controls. Black, total cells (∼5,000/sample); red, mCherry-positive invaded cells (∼500/sample). Dots indicate biological replicates; bars indicate means (*n* = 3). **(e)** Distributions of NF-κB reporter activity across infection time points, measured by GFP MFI, in mCherry-positive invaded (red) and uninvaded (gray) cells. Representative of three biological replicates. **(f)** Effects of NEMO loss and complementation on intracellular CAB2 replication in HeLa cells. Top, Western blot analysis of control and NEMO-knockout (NEMO^−/−^) HeLa cells (GAPDH, loading control). Bottom, intracellular bacterial loads over time (as in Fig. 4b). Mean ± s.d. (*n* = 3). Student’s *t*-test versus control at each time point: ns, *P* ≥ 0.05; *, *P* < 0.05; **, *P* < 0.01; ***, *P* < 0.001. **(g)** NF-κB reporter activity in mock- or CAB2-infected Caco-2 cells (4 h), treated with DMSO, BMS-345541 (10 μM), or TNF-α (1 μg/mL). Representative of three biological replicates (∼20,000 cells/sample). **(h)** Intracellular CAB2 loads in Caco-2 cells treated as in **g**, plus untreated controls. Mean ± s.d. (*n* = 3); statistics as in **f**.

To validate NF-κB activation independently of the RNA-seq data, we used a HeLa reporter cell line carrying a genome-integrated GFP reporter driven by NF-κB-responsive elements (**Methods**). Although NF-κB activation was inferred from RNA-seq data collected in Caco-2 cells, we used HeLa cells for functional validation because knockout lines were available in this background, and HeLa cells are also a well-established model of *V. para* infection in our laboratory^8,16^. Flow cytometry analysis showed that NF-κB-induced GFP signal progressively accumulated over the course of infection, consistent sustained NF-κB activation (**Fig. 6d**). This fluorescence-based reporter readout also provides independent support for the NF-κB activation inferred from the RNA-seq data. Because canonical NF-κB activation depends on the IKK complex, in which NEMO functions as an essential regulatory component, we next tested this response in NEMO knockout cells. In contrast to wild-type cells, NEMO knockout cells showed no significant increase in reporter signal over time, indicating that infection-induced NF-κB activation requires NEMO-dependent canonical signaling (**Fig. 6d**).

We next asked whether NF-κB activation occurred preferentially in host cells containing intracellular bacteria. To test this association, we used FACS to distinguish invaded cells based on bacterial fluorescence. Across all time points, invaded cells showed stronger NF-κB activation than the rest of the population (**Fig. 6d, e**), confirming that NF-κB activation is enriched in cells that contain intracellular *V. para*. The low level of detectable activation in bystander cells may reflect the spread of inflammatory signaling through secondary signals released from infected cells.

Finally, we asked whether NF-κB activation simply reflects the host response to infection, or whether it influences the course of intracellular bacterial replication. We quantified intracellular bacterial load over the infection time course in control HeLa cells and NEMO knockout cells. Recovered bacterial counts were significantly reduced in NEMO knockout cells compared with control cells (**Fig. 6f**). Consistent with this genetic evidence, pharmacological inhibition of IKK signaling produced a similar outcome in Caco-2 cells. Treatment with the IKK inhibitor BMS-345541^48^, at a concentration that did not reduce the viability of either Caco-2 cells or *V. para* (**Fig. S4**), suppressed infection-induced NF-κB reporter activation (**Fig. 6g**). Under this condition, intracellular bacterial replication was significantly impaired during infection (**Fig. 6h**). Conversely, acutely enhancing NF-κB activation by adding TNF-α during bacterial invasion also reduced intracellular bacterial amount (**Fig. 6g, h**). Thus, both loss and excessive activation of NF-κB signaling were unfavorable for intracellular *V. para* replication.

Together, these results indicate that *V. para*-induced NF-κB activation is not simply a marker of the host inflammatory response but instead reflects a host cell state that supports efficient intracellular bacterial replication. Both reduced and excessive NF-κB signaling impaired intracellular bacterial survival and replication, suggesting that *V. para* depends on an optimal range of NF-κB activity during infection, consistent with a *Goldilocks Principle*, in which neither insufficient nor excessive signaling is favorable. Thus, *V. para* exploits a finely tuned host NF-κB response to create an intracellular environment permissive for bacterial expansion.

The manipulation of NF-κB activity by *V. para* is conceptually consistent with macrophage studies using other intracellular pathogens such as *Legionella pneumophila* and *Mycobacterium tuberculosis*, where NF-κB-associated survival signaling can delay or restrain caspase-3 linked apoptosis and thereby support intracellular bacterial survival and expansion^49,50^. However, *V. para* manipulation of NF-κB activity contrasts the regulation of NF-κB activity by the extracellular pathogen *Yersinia spp.* encoding the acetyltransferase YopJ/P that inhibits all MAPK pathways and the NF-κB pathway^51^. Interestingly, *V. para* encodes a homologue of YopJ/P that is also an acetyltransferase but is only able to inhibit MAPK pathways and not NF-κB activity^52^.

Although Caco-2 cells are widely used to model intestinal epithelial infections ^53–55^, their cancer-derived origin and conventional culture conditions may not fully recapitulate the metabolic properties^56^ and inflammasome expression^57^ of primary intestinal epithelial cells. Similarly, the use of the *V. para* CAB2 strain minimizes cytotoxicity and enables detailed investigation of invasion and intracellular replication. However, this strain excludes potential hemolysin-mediated effects on host cells^58^ as well as the role of potential regulatory interactions between T3SS1 and T3SS2 in host-cell invasion, both of which also remain beyond the scope of this study^59^.

### Metabolic implications of *V. para* nutrient acquisition during intracellular infection

A central feature of bacterial adaptation was the temporal remodeling of nutrient acquisition, which provided clues to the intracellular niche environment encountered by *V. para* at different stages of intracellular infection. Notably, genes involved in sulfur-containing amino acid biosynthesis and transport showed an early response at 1.5 h, preceding the broader activation of amino acid acquisition systems (**Fig. 3b**; **Fig. S5a**). Sulfate assimilation genes were also induced at 1.5 h (**Fig. S5b**), suggesting a potential role for sulfur metabolism in early intracellular adaptation beyond protein synthesis. Sulfur metabolism was also enriched in cluster 6, which showed sustained transcriptional activation throughout infection (**Fig. 2c**). Similar sulfur-related metabolic adaptations have been reported in *E. coli* under sulfur-limiting conditions^60^ and in *V. fischeri* during host symbiosis^61^. Meanwhile, host sulfur amino acid metabolism showed increasingly negative enrichment during infection (**Fig. S6a**). These contrasting responses raise the possibility that both the host and bacterium face limited sulfur nutrient availability during intracellular infection, with *V. para* enhancing sulfur acquisition and biosynthesis while host cells reduce sulfur catabolism to conserve these resources. In addition, glutathione biosynthesis genes of *V. para* were elevated relative to the pre-invasion MEM condition (**Fig. S5c**), suggesting a possible contribution of sulfur metabolism to redox homeostasis following host-cell entry.

Beyond the amino acid pathways discussed in the study, carbon source transport showed particularly clear temporal dynamics. Gluconate and hexosephosphate transporters were induced at the earliest intracellular time point. Although expressed at relatively low levels, activation of a UhpT-like hexosephosphate transporter may mark the transition to the host cytosol, as reported in *Salmonella*^62^ and *Listeria*^39^. During rapid intracellular proliferation, broader induction of transporters for sugars and other glycolysis related substrates, including multiple phosphotransferase system systems, likely reflects increased carbohydrate demand and may enable the co-utilization of several carbon sources^38,63^. Consistent with this broader expansion of carbon acquisition, the concurrent induction of fatty acid degradation at 4 h suggests that *V. para* may utilize fatty acid-derived carbon to supplement carbohydrate utilization for replication (**Fig. 2c**). More broadly, fatty acid-derived metabolites can directly influence carbohydrate metabolic flux^64^, highlighting the potential coordination between lipid and carbohydrate metabolism under the high metabolic demands of rapid growth. Together, these findings suggest that the rapid replication phase is supported by broad and coordinated carbon utilization. As infection progressed to later stages, the carbon-acquisition program shifted toward systems associated with organic acids and metabolic intermediates. Increased transport of succinate and other organic acids may reflect access to host TCA cycle intermediates. Formate and acetate associated systems could additionally reflect the accumulation, export or recycling of bacterial fermentation products^65,66^.

In contrast to the stage-specific dynamics of sugar related transporters, iron acquisition systems displayed a broader and more sustained response. Many genes in the Fur regulon, including those involved in TonB2-dependent transport, ferric iron uptake, and several siderophore transport systems, peaked during pre-invasion incubation in MEM (**Fig. S2a**). This response may reflect increased iron demand following the transition to serum-free MEM, which lacks supplemental iron, rather than an infection-specific response. Although vibrioferrin uptake genes peaked at 1.5 h, genes involved in its biosynthesis and export were strongly induced from 4 h onward (**Fig. S2b**). This pattern coincided with the progressive induction of RyhB (**Fig. S2c**), a small RNA previously shown to promote vibrioferrin biosynthesis by stabilizing *pvs* operon transcripts^67^. This suggests that RyhB-mediated regulation may enhance vibrioferrin production during intracellular growth, potentially facilitating the utilization of host-associated iron^68^. Following a similar temporal pattern, heme uptake and utilization genes were preferentially induced at 4 and 6.5 h (**Fig. S2d**), suggesting that host-derived heme may serve as another potential iron source. Together, these findings suggest that *V. para* flexibly adapts its iron acquisition strategies to changes in iron availability and sources, both during transitions between culture media and upon entry into host cells.

### Physiological implications of host transcriptional remodeling during infection

Amino acid metabolism offered a further view of how bacterial nutrient adaptation may track changes in host-cell metabolism. At 1.5 h post-infection, infected host cells showed enriched mTORC1 signaling and oxidative phosphorylation, indicating an early metabolically active state with anabolic signaling and mitochondrial respiration (**Fig. 6a**; **Fig. S6b**). This state was not maintained as infection progressed. At later time points, mTORC1 and oxidative phosphorylation signatures declined, while unfolded protein response pathways increased and solute carrier-mediated amino acid transport became progressively depleted (**Fig. S6c**). In addition, the GCN2-mediated amino acid deficiency response showed modest positive enrichment early in infection but declined thereafter (**Fig. S6c**), suggesting a transient response to amino acid limitation rather than sustained starvation. Together, these changes suggest a transition from early anabolic and respiratory activity toward a more stress-associated metabolic state, accompanied by altered amino acid homeostasis. In parallel, intracellular *V. para* induced broad amino acid transport and biosynthetic programs, including shikimate derived aromatic amino acid biosynthesis (**Fig. 3**; **Fig. 4a**). This temporal alignment suggests that *V. para* increases its capacity for amino acid uptake and synthesis as the host cell becomes metabolically stressed and potentially more nutrient limited.

Beyond these pathway-specific metabolic changes, the host transcriptome exhibited a broader temporal organization across infection. Although transcriptional remodeling in the host was less extensive than that observed in *V. para* (**Fig. 1e**), principal component analysis clearly separated transcriptional patterns in invaded cells from mock controls and further resolved the progression of infection (**Fig. 1d**). PC2 primarily distinguished invaded cells from mock controls, with genes contributing to PC2 enriched in multiple host pathways, including TNF-α signaling via NF-κB and inflammatory response (**Fig. S7a**). These pathways recapitulated the infection-associated signatures identified by whole-transcriptome GSEA (**Fig. 6a**). PC1, meanwhile, ordered the infected samples from 1.5 h, 4 h, and 6.5 h and therefore appeared to resolve the temporal remodeling of infected host cells across the intracellular infection trajectory (**Fig. 1d**).

Analysis of genes contributing to PC1 revealed two opposing temporal patterns in host pathway activity. First, compared with uninfected mock cells, genes associated with apoptosis, MYC-linked growth control and EMT/TGF-β-related epithelial remodeling were strongly induced at 1.5 h, returned to near-mock levels by 4 h and declined further at 6.5 h (**Fig. S7b**). These coordinated changes indicate a rapid early host response that is progressively attenuated as infection proceeds. In contrast, genes associated with heme and bile acid metabolism were initially repressed and then increased toward late infection (**Fig. S7c**). These changes may reflect progressive remodeling of host iron and redox homeostasis, as well as lipid and sterol metabolism, as the intracellular bacterial burden increases. The latter is consistent with the increasing expression of a bacterial lipase during intracellular infection (**Fig. 5a**) and its previously reported effects on host lipid homeostasis^12^. Together, these PC1-associated trajectories outline a temporal progression of the host response during intracellular *V. para* infection. Together, these opposing trajectories place the 4 h time point as a pivotal transition between the early invasion-associated response and the later emergence of metabolic stress. Notably, the relatively near-mock host profile at 4 h coincides with extensive intracellular bacterial replication, supporting the possibility that this transitional host state provides a particularly permissive environment for bacterial growth.

In summary, in this study we revealed a temporal definition of transcriptomic changes in both bacteria and host cells during the intracellular infection cycle of *V. para* using dual RNA-seq. After host-cell entry, *V. para* underwent ordered transcriptional transitions, including broad remodeling of nutrient acquisition and metabolism during intracellular replication, followed by late induction of secretion and flagellar associated programs. Among these metabolic responses, methionine metabolism was activated upon initial infection while arginine and aromatic amino acid biosynthesis was activated during mid-stage infection, and bacterial production of aromatic amino acids was required for efficient intracellular replication. Analysis of the host response further revealed an unexpected relationship between inflammatory signaling and bacterial fitness: rather than acting as a uniformly restrictive response, moderate NF-κB signaling was beneficial to intracellular bacterial replication. Overall, we observe the pathogenic *V. para* subtly entering and adapting to a new intracellular environment without setting off major alarms. This strategy is consistent with previous observations showing *V. para* is coated in a hepta-acylated LPS and sheathed flagellum^69^ that weakly activates host immune signaling^13^. Together, these findings show that intracellular infection depends not only on bacterial metabolic adaptation, but also on the establishment of a finely balanced host-cell state that supports bacterial expansion. Future studies will be directed at understanding how *V. para* engages these host signaling pathways to establish its intracellular niche.

## Methods

### Bacteria strains and cell culture

The invasive *V. para* strain CAB2, previously described^8^, was derived from POR1 by deletion of *exsA*, which encodes the transcriptional activator of T3SS1. POR1 was derived from the clinical isolate RIMD2210633 by deletion of the TDH-encoding genes^70,71^. *V. para* strains were cultured at 30°C in marine LB (MLB), consisting of Luria–Bertani (LB) medium (Fisher, BP97235) supplemented with NaCl to a final concentration of 3% (w/v), with the indicated antibiotics when required. The *E. coli* SM10 λpir strain (Genprice, S0049) was cultured in LB medium containing the indicated antibiotics when required.

HeLa and Caco-2 cells were obtained from ATCC. Caco-2 cells were routinely cultured in MEM/EBSS (Cytiva, SH30024) supplemented with 20% FBS (Sigma, F2442) and 1% penicillin–streptomycin (Sigma, P0781). HeLa cells were routinely cultured in high-glucose DMEM (Sigma, D5796) supplemented with 10% FBS, 1% penicillin–streptomycin and 1 mM sodium pyruvate (Gibco, 11360070).

### *V. para* infection and gentamicin protection assay

All infection experiments were performed using a gentamicin protection assay. Overnight cultures of *V. para* were diluted to an OD₆₀₀ of 0.3, supplemented with 100 μM TDC (Sigma, T0875) to induce T3SS2 expression, and cultured at 37°C for 90 min. Infection were performed in cell culture medium without FBS or antibiotics.

Following induction, bacteria were diluted in cell culture medium to prepare the infection medium at a multiplicity of infection (MOI) of 10. To synchronize infection, the plates were centrifuged at 1,000 rpm for 5 min and subsequently incubated at 37°C with 5% CO₂ for 90 min to allow bacterial invasion. The infection medium was then removed, and the cells were washed with PBS to remove extracellular bacteria. Fresh medium containing 100 μg/mL gentamicin (Thermo, 455310050) was added to eliminate the remaining extracellular *V. para*. At the indicated time points, infected cells were collected for colony-forming unit (CFU) assay or FACS. Mock-infected cells were processed identically, except that no bacteria were added to the infection medium.

For the intracellular CFU assay, HeLa or Caco-2 cells were seeded in 24-well plates. At indicated time points after gentamicin addition, cells were washed with PBS and then lysed in 1 mL of 0.5% Triton X-100 (Fisher, BP151) in PBS. Plates were shaken at 100 rpm for 10 min, followed by vigorous pipetting to ensure complete cell lysis. Cell lysates were serially diluted 10-fold, and 10 μL of each dilution was spotted onto MLB agar plates. Plates were incubated overnight at 30°C, and colonies were counted to determine CFU.

For pharmacological treatments, BMS-345541 (MCE, HY-10519) was added during the 90-min invasion period and maintained in the gentamicin-containing medium throughout the remainder of the infection. Purified TNF-α, prepared as previously described^72^, was added only during the 90-min invasion period.

### Construction of *V. para* deletion mutants and complemented strains

The suicide vector pDM4 was used to generate gene deletions in CAB2. Briefly, 0.8∼1.0-kb regions upstream and downstream of each target gene were amplified with Phusion DNA polymerase (Thermo, F530L) from genomic DNA extracted using the EZ-10 Spin Column Bacterial Genomic DNA Mini-Preps Kit (Bio Basic, BS624). The upstream and downstream fragments were designed with overlapping ends and fused by overlap PCR. The resulting fragment was digested with SmaI and XhoI (Thermo, FD0664 and FD0694), ligated into pDM4 digested with the same enzymes using T4 DNA ligase (Thermo, EL0011), and transformed into *E. coli* SM10 λpir strain by heat shock. Transformants were selected on LB agar containing 25 μg/mL chloramphenicol. SM10 competent cells were prepared by incubating log-phase bacteria cells in 0.1 M CaCl₂ on ice for 1 h. SM10 carrying the recombinant plasmid was conjugated with CAB2 by mixing the two strains at a 1:1 ratio and incubating the mixture on an MLB agar plate. Conjugants were selected on MLB agar containing 25 μg/mL chloramphenicol (Fisher, BP904). To remove the *sacB*-containing pDM4 plasmid, conjugants were counter-selected on MLB agar containing 15% (w/v) sucrose. Individual sucrose-resistant colonies were screened by colony PCR (Thermo, K1082) across the target locus, and successful deletions were confirmed by Sanger sequencing.

For construction of the complemented strains, the coding sequences of *aroA* and *aroB* were cloned into pBAD-Fix between the SacI and SalI sites. The resulting plasmids were transformed into SM10 and introduced into the corresponding CAB2 Δ*aroA* or Δ*aroB* mutant by conjugation. Transconjugants were selected on MMM agar containing 250 μg/mL kanamycin (Fisher, BP906). During gentamicin protection assays, expression of the complemented genes was induced with 0.02% arabinose (w/v) during TDC induction in MLB containing kanamycin.

Primers used for *V. para* strain and complemental plasmid construction are listed in **Table S4**.

### Generation of NEMO knockout and NF-κB reporter cell lines

For generation of NEMO knockout (KO) cells, sgRNAs targeting NEMO (5′-GGCAGAGCAACCAGATTCTG-3′) was cloned into the lentiCRISPR v2 vector (Addgene plasmid #98290). The NF-κB reporter construct was generated from pCDH-CMV-MCS-EF1α-Hygro (System Biosciences, CD515B-1). The original CMV promoter upstream of EGFP was replaced with seven tandem κB response elements fused to a minimal IFN-β promoter.

Lentiviruses were produced by co-transfecting 293T cells with the indicated lentiviral construct, psPAX2, and pMD2.G. Viral supernatants were collected, filtered through a 0.45-μm filter, and used to transduce the indicated cells. Transduced cells were selected with the appropriate antibiotics to generate stable cell lines.

For the NF-κB reporter cell line, transduced HeLa or Caco-2 cells were selected with the appropriate antibiotic and maintained as stable polyclonal populations. For NEMO KO, HeLa STING^−/−^ cells were used because endogenous STING expression is low in HeLa cells and its loss is not expected to substantially alter the parental phenotype. Cells were transduced with the NEMO-targeting lentivirus, followed by single-cell cloning. KO clones were validated by western blotting, functional assays, and Sanger sequencing.

### Flow cytometry and cell sorting

To collect infected cells for dual RNA-seq, Caco-2 cells were seeded in 6-well plates and infected with CAB2 carrying pMW-GFP at an MOI of 10. At 1.5, 4.0, and 6.5 h after gentamicin addition, cells were detached with trypsin and collected by centrifugation at 500 × *g* for 5 min. Cell pellets were resuspended in FACS buffer containing 1% FBS and 1 mM EDTA in PBS and kept on ice until sorting. Cell suspensions were passed through a 35-μm cell strainer (Falcon, 352235) and sorted using a five-laser BD FACSAria II SORP (BD Biosciences). Both the sample holder and collection tube rack were maintained at 4°C. Cells were first gated by forward and side scatter to select the major cell population and exclude debris, followed by SSC-A versus SSC-H gating to select singlets. Infected cells were then identified based on GFP fluorescence in the FITC channel relative to cellular autofluorescence in the APC channel. GFP-positive cells were sorted into 1.5-mL tubes containing PBS with 2% FBS and kept on ice. Mock-infected cells were sorted using the same gating strategy, except that the final GFP-positive gate was omitted. Approximately 1 × 10⁵ cells were collected from each fraction for RNA isolation, with two biological replicates collected per condition.

For NF-κB activation measurements, HeLa or Caco-2 NF-κB reporter cells were infected with CAB2 carrying pMW-mCherry at an MOI of 10. At the indicated times after gentamicin addition, cells were harvested, processed, and gated for single cells as described above. Infected cells were identified by mCherry fluorescence in the PE channel, and NF-κB activity was measured by EGFP fluorescence in the FITC channel.

### Dual RNA-seq sample preparation and total RNA extraction

For sorted infected and mock-infected host-cell samples, sorted cells were pelleted in 1.5-mL tubes by centrifugation at 800 × *g* for 5 min. Each pellet was resuspended directly in 0.5 mL TRIzol reagent (Invitrogen, 15596026), and RNA was extracted according to the manufacturer’s instructions. During isopropanol precipitation, 1 μL GlycoBlue (Invitrogen, AM9515) was added to each sample, and the samples were incubated at −20°C overnight. RNA was pelleted by centrifugation at 12,000 × *g* for 15 min at 4°C. The supernatant was carefully removed, and the RNA pellet was air-dried and dissolved in 20 μL nuclease-free water (Invitrogen, AM9932).

For *V. para* control samples, two biological replicates were prepared for each condition. For the TDC control, 50 μL of CAB2 culture was collected after 90 min of TDC induction. For the MEM control, bacteria collected after TDC induction were resuspended in 2 mL MEM and added to 6-well plates. The plates were centrifuged under the same conditions used to synchronize infection and incubated at 37°C with 5% CO₂ for 90 min. Bacteria from both control conditions were collected by centrifugation at 10,000 × *g* for 1 min. Bacterial pellets were resuspended in 50 μL freshly prepared lysis buffer containing 15 mg/mL lysozyme (Sigma, SAE0152) in TE buffer (Invitrogen, AM9858) and 3.5 mg/mL proteinase K (QIAGEN, 19131). Samples were incubated at 37°C with shaking for 15 min. Bacterial RNA was purified using 100 μL of RNAClean XP beads (Beckman, A63987) according to the manufacturer’s instructions and eluted in 20 μL nuclease-free water.

RNA concentration was measured using the Qubit RNA High Sensitivity Assay Kit (Invitrogen, Q32852).

### Library construction

Library construction was adapted from previously described protocols and further customized for this study^27,28^. All reactions were assembled in 0.2-mL PCR tubes using master mixes prepared from the shared reaction components.

Briefly, genomic DNA was removed by treating RNA samples with DNase I (NEB, M0303S) in the presence of SUPERase·In RNase inhibitor (Invitrogen, AM2694) at 25°C for 30 min. RNA was then purified using 1.8× volumes of RNAClean XP beads and eluted in 11 μL nuclease-free water. Purified RNA was subjected to RNase H-based rRNA depletion^28^. For each reaction, 200–400 ng RNA in 10 μL was used in a final reaction volume of 20 μL. For host-cell-derived samples, rRNA depletion was performed using the NEBNext rRNA Depletion Kit v2 (NEB, E7400) according to the manufacturer’s instructions, with the addition of 1 μL customized *V. para* rRNA depletion probes (100 μM; sequences listed in **Table S5**) to the hybridization reaction. For *V. para* control samples, the hybridization reaction contained 10 μL RNA, 3 μL customized *V. para* rRNA depletion probes (100 μM), 1 μL 1 M Tris-HCl, pH 7.5 (Invitrogen, 15567027), 0.6 μL 5 M NaCl (Invitrogen, AM9760G), and 5 μL nuclease-free water. Following probe hybridization, RNase H digestion, and DNase I digestion, RNA was purified using 1.8× volumes of RNAClean XP beads and eluted in 5 μL nuclease-free water.

First-strand cDNA synthesis and template switching were adapted from the SMART-seq2 framework^73^. For the two biological replicates of each condition, reverse transcription was performed using random hexamer primers containing distinct 5′ constant sequences that served as replicate-specific indices (**Table S5**). For each reaction, 5 μL RNA was combined with 1 μL 10 μM reverse-transcription primer, 1 μL 10 mM dNTPs, and 1.5 μL nuclease-free water.

Samples were mixed by pipetting, incubated at 65°C for 5 min, immediately placed on ice, and maintained on ice for at least 1 min. An 11-μL reverse-transcription (RT) mixture containing 4 μL 5× SuperScript IV buffer, 1 μL 100 mM DTT, 1 μL 10 μM template-switching oligonucleotide, 3 μL nuclease-free water, 1 μL SUPERase·In RNase inhibitor, and 1 μL SuperScript IV reverse transcriptase (Invitrogen, 18090050) was added to each denatured RNA sample on ice. Reactions were mixed by pipetting and incubated at 23°C for 10 min, 50°C for 45 min, and 80°C for 10 min, followed by a hold at 4°C.

Following RT, second-strand synthesis and cDNA amplification were performed by PCR. Each 25-μL reaction contained 10 μL of the RT product, 0.5 μL each of 10 μM Adaptor_primer and ISPCR_primer, 12.5 μL 2× KAPA HiFi HotStart ReadyMix (Roche, KK2601), and 1.5 μL nuclease-free water. PCR was performed with the following program: 98°C for 3 min; 9 cycles of 98°C for 20 s, 64°C for 20 s, and 72°C for 3 min; followed by 72°C for 5 min and a hold at 4°C. The amplified cDNA was purified using 1.0× volumes of AMPure XP beads and eluted in 10 μL nuclease-free water. DNA concentration was measured using the Qubit 1× dsDNA High Sensitivity Assay Kit (Invitrogen, Q33230).

For each condition, 5 ng cDNA from each of the two biological replicates was pooled to obtain 10 ng total cDNA for tagmentation using the TruePrep Index Kit V2 for Illumina (Vazyme, TD501). Tagmentation was performed according to the manufacturer’s instructions, except that the transposase was diluted and titrated to produce libraries ranging from 250 to 600 bp after PCR amplification. Libraries were amplified using primers containing P5 or P7 adapters and condition-specific index combinations (**Table S6**). Libraries were pooled for sequencing at ratios designed to achieve target depths of 10 Gb per infected-cell replicate, 5 Gb per mock-infected host-cell replicate, and 2 Gb per *V. para* control replicate. Sequencing was performed by Novogene on an Illumina NovaSeq X platform with 150-bp paired-end reads.

All primers and oligonucleotides used for library construction are listed in **Table S6**.

### Dual RNA-seq data processing and read quantification

Raw paired-end sequencing reads were first quality-filtered using Cutadapt^74^. Bases with Phred quality scores below 10 were trimmed from both ends, and read pairs containing more than three ambiguous bases or a read shorter than 100 nt after trimming were discarded.

Two biological replicates pooled within each library were demultiplexed using replicate-specific sequences introduced during reverse transcription. Read 1 sequences beginning with GNGTGAA or GNCACAA were assigned to replicate 1 or replicate 2, respectively. The transposase-related sequence and the template-switching-related sequence were trimmed from the 3′ end, and reads shorter than 20 nt after trimming were discarded. Separate Cutadapt runs were performed for the two replicates using the following settings: -g ^GNGTGAA or -g ^GNCACAA, --no-indels, --minimum-length 20, --match-read-wildcards, --discard-untrimmed, -a CTGTCTCTTATACACATCTC, and -a CCCATGTACTCTGCGTTGAT. Owing to the customized library architecture, only the processed Read 1 sequences were retained and analyzed as single-end reads in all downstream workflows.

For host transcriptome analysis, all host-cell-derived samples, including infected and mock-infected samples, were aligned to the human GRCh38 reference genome using STAR^75^. Demultiplexed reads of at least 50 nt were retained for alignment, and alignments were output as coordinate-sorted BAM files using --outSAMtype BAM SortedByCoordinate. Gene-level counts were generated using featureCounts^76^ with the Ensembl release 113 annotation filtered to chromosomes 1–22, X, Y, and MT. featureCounts was run with the following settings: -s 2 -t gene -g gene_id. TPM values were calculated from the resulting gene-count table after excluding mitochondrial genes, and the TPM table was used for downstream analyses.

For *V. para* transcriptome analysis, reads from infected host-cell samples and *V. para* control samples were aligned to the *V. para* RIMD2210633 reference genome (GCF_000196095.1) using Bowtie2^77^. Alignments were converted from SAM to BAM format and sorted using SAMtools^78^. Gene-level counts were generated using featureCounts^76^ with the corresponding genome annotation (GCF_000196095.1_ASM19609v1) and the following settings: -s 2 -t gene -g locus_tag. TPM values were calculated after excluding rRNA and ssrA genes from both the reported gene set and the normalization denominator, and the resulting TPM table was used for downstream analyses.

Raw sequencing reads in FASTQ format and processed gene-count data have been deposited in the NCBI Gene Expression Omnibus under accession **GSE341606**.

### Temporal clustering and functional enrichment analysis of *V. para* gene expression

*V. para* gene expression was quantified as transcripts per million (TPM) using featureCounts-derived read counts and annotated gene lengths. Genes encoding rRNAs and *ssrA* were excluded before TPM calculation. For temporal expression clustering, genes with a summed TPM of at least 5 across the 10 samples representing five conditions were retained.

For each gene, TPM values were divided by the mean TPM across the selected samples and transformed as log₂(normalized expression + 1). Biological replicates were then averaged for each condition. K-means clustering was performed on the resulting condition-level expression profiles using scikit-learn with *k* = 7 and k-means++ initialization. The analysis was repeated with 100 random seeds, and the solution with the lowest within-cluster sum of squares (inertia) was retained. To visualize the relationships among gene-expression trajectories, the same normalized expression profiles were projected into two dimensions using Uniform Manifold Approximation and Projection (UMAP)^79^, with genes colored according to their K-means cluster assignments. UMAP was used only for visualization and did not contribute to cluster assignment.

Functional enrichment analysis was performed separately for genes in each K-means cluster. An integrated functional reference was assembled by combining KEGG pathways (code vpa)^29^, predicted regulons from RegPrecise^31^, and manually curated gene sets. Gene Ontology (GO) Biological Process annotations were processed using GMT Helper^80^. GO enrichment was performed separately to prevent the large number of overlapping GO terms from influencing the enrichment statistics of other functional gene sets. Enrichment was assessed using hypergeometric tests with the enricher function in clusterProfiler^81^. Gene sets containing 3–200 genes were tested in both analyses. Functional terms with nominal *P* values < 0.05 were retained without multiple-testing correction, whereas GO terms were retained based on Benjamini–Hochberg-adjusted *P* values < 0.05.

### Host differential expression and gene set enrichment analysis (GSEA)

For host differential expression analysis, raw gene counts were analyzed using DESeq2^82^. Each comparison was performed independently using a design based on experimental condition. Genes with fewer than 10 total counts across the samples included in each comparison were excluded before analysis. Differential expression was assessed between infected and mock-infected cells at 1.5, 4.0, and 6.5 h after gentamicin addition.

For gene set enrichment analysis (GSEA), genes from each infected-versus-mock comparison were ranked by the DESeq2 Wald test statistic. Ensembl gene identifiers were converted to gene symbols using org.Hs.eg.db. Genes without a mapped symbol or DESeq2 statistic were excluded, and when multiple Ensembl identifiers mapped to the same gene symbol, the entry with the largest absolute Wald statistic was retained. Pre-ranked GSEA was performed separately for each infection time point using the GSEA function in clusterProfiler^81^ and the Hallmark gene sets obtained from MSigDB using msigdbr^47,83^. All successfully evaluated gene sets were retained in the initial GSEA output (*P*-value cutoff set to 1). Statistical significance was subsequently assessed using the nominal *P* value and false discovery rate-adjusted *P* value. Normalized enrichment scores and running enrichment-score profiles were used for downstream visualization.

For selected pathways, running enrichment-score curves and the positions of gene-set members within the ranked gene lists were extracted using enrichplot and plotted across the three infection time points.

## Data, Code, and Materials Availability

All data and code needed to evaluate and reproduce the results in the paper are present in the paper and/or the Supplementary Materials. All data and code needed to evaluate and reproduce the results in the paper are present in the paper and/or the Supplementary Materials. This study did not generate new materials. Raw sequencing reads (FASTQ) and processed gene-count data generated in this study have been deposited in the NCBI Gene Expression Omnibus (GEO) under accession number GSE341606. The dataset will be made publicly available upon publication.

## Author Contributions

Conceptualization: Y.Z., K.O., L.N.K., C.S.

Methodology: Y.Z., C.S.

Experimentation: Y.Z., C.S., S.V., J.J.

Data Analysis: Y.Z., K.O., L.N.K., C.S.

Resources: K.O., Z.J.C.

Writing original manuscript: Y.Z.

Editing manuscript: Y.Z., K.O., L.N.K., S.V., C.S.

## Acknowledgements

We thank the Orth lab and Ping Shen for productive discussions and editing. The study was funded by the Welch Foundation I-1561 (KO), Once Upon a Time Foundation (KO) and an R01CA299257 (ZJC). KO is a W.W. Caruth, Jr. Biomedical Scholar with an Earl A. Forsythe Chair in Biomedical Science. ZJC is a George L. MacGregor Distinguished Chair in Biomedical Sciences.

## Supplementary Materials

Figs. S1 to S7

Tables S1 to S6

