## Supplementary material for "*Vibrio parahaemolyticus* metabolically adapts while tempering the host immune response during host cell invasion": Supp Data

### Supplementary Figures

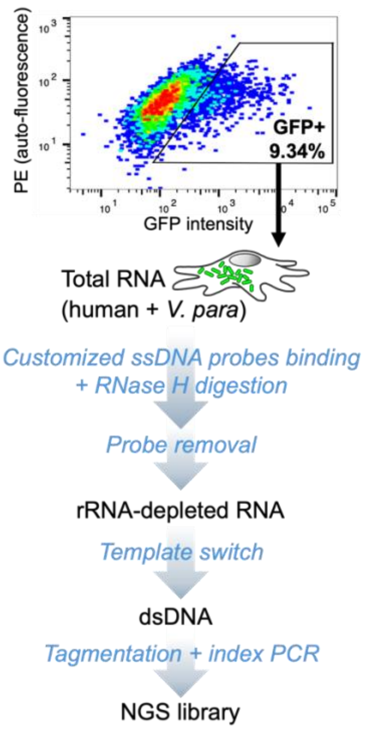


**Figure S1 Dual RNA-seq library construction workflow.**

Total RNA was extracted from infected cells. Human and *V. para* rRNAs were hybridized to a combined pool of host- and pathogen-specific ssDNA probes and depleted by RNase H digestion. The rRNA-depleted transcripts were then subjected to first- and second-strand cDNA synthesis using a template-switching reaction, followed by library indexing and fragmentation through tagmentation.

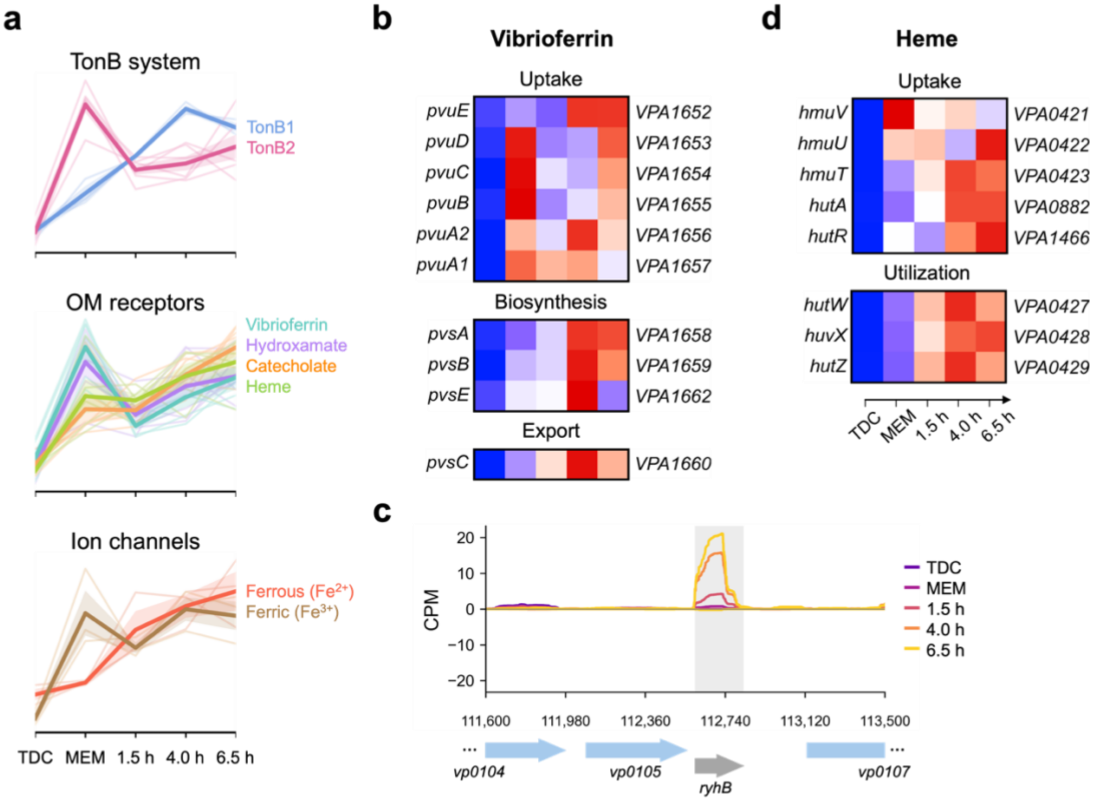


**Figure S2 Expression dynamics of *V. para* iron acquisition systems.**

**(a)** *Z*-score-normalized expression of iron acquisition genes grouped into three categories: TonB systems, outer membrane (OM) receptors, and ion transporters. Thin lines represent individual genes, and thick colored lines indicate group means. Thin lines show individual gene expression averaged across two biological replicates; thick colored lines represent group means.

**(b)** *Z*-score-normalized expression of genes involved in vibrioferrin uptake, biosynthesis, and export. Heatmaps show mean expression across two biological replicates. Rows represent individual genes, with gene names shown on the left and locus tags on the right.

**(c)** RNA-seq coverage profiles of the RyhB genomic region across conditions. Coverage is shown as counts per million (CPM) for both strands, averaged across two biological replicates. Genomic coordinates and gene annotations are indicated along the x-axis. Conditions are distinguished by color.

**(d)** *Z*-score-normalized expression of genes involved in heme uptake and utilization, displayed as in **b**.

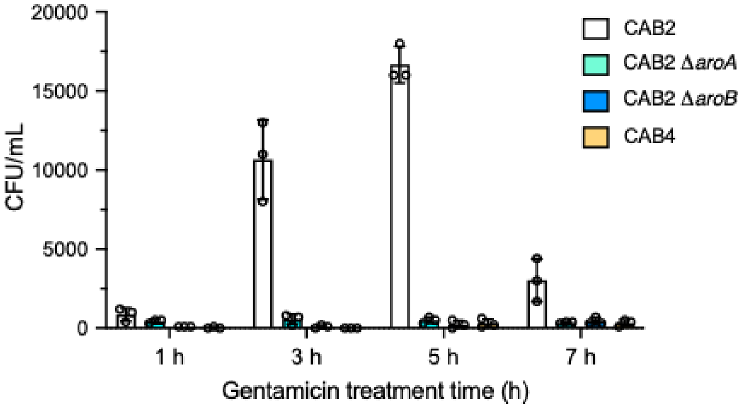


**Figure S3 Intracellular replication of *V. para* strains in HeLa cells.**

Intracellular replication of *V. para* CAB2, Δ*aroA*, Δ*aroB*, and the invasion-defective strain CAB4 in HeLa cells. Cells were infected for 90 min and then incubated in gentamicin-containing medium. Intracellular bacterial loads were determined at the indicated time points post-infection. Data are shown as mean ± s.d. ($n$ = 3 biological replicates).

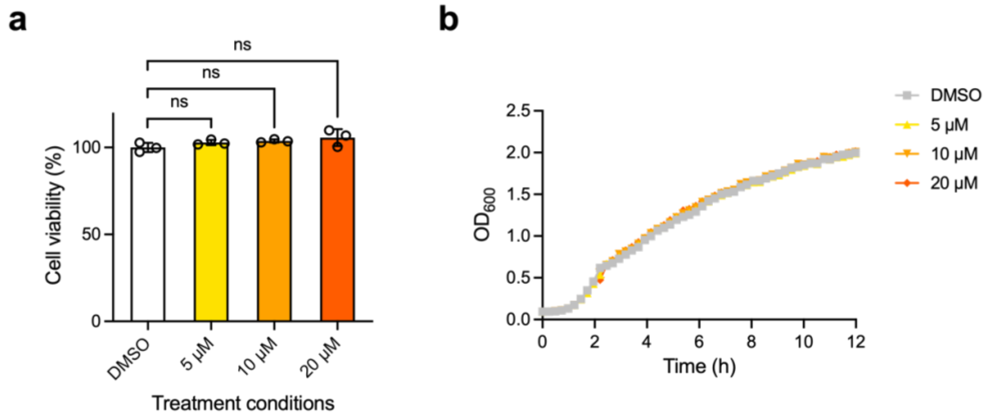


**Figure S4 BMS-345541 treatment does not impair Caco-2 cell viability or *V. para* growth at the concentrations tested.**

**(a)** Relative viability of Caco-2 cells treated with 5, 10, or 20 µM BMS-345541 for 7 h, corresponding to the duration of the infection assay. Viability was normalized to that of the DMSO-treated control. Data are shown as mean ± s.d. ($n$ = 3 biological replicates; unpaired *t*-tests, ns, $P > 0.05$).

**(b)** Growth of *V. para* CAB2 in MLB containing 5, 10, or 20 µM BMS-345541 at 37°C. Overnight cultures were diluted to an initial $OD_{600}$ of 0.02 in 96-well plates, and growth was monitored by measuring $OD_{600}$ every 15 min for 12 h using a plate reader. Data are shown as mean ± s.d ($n$ = 4 biological replicates).

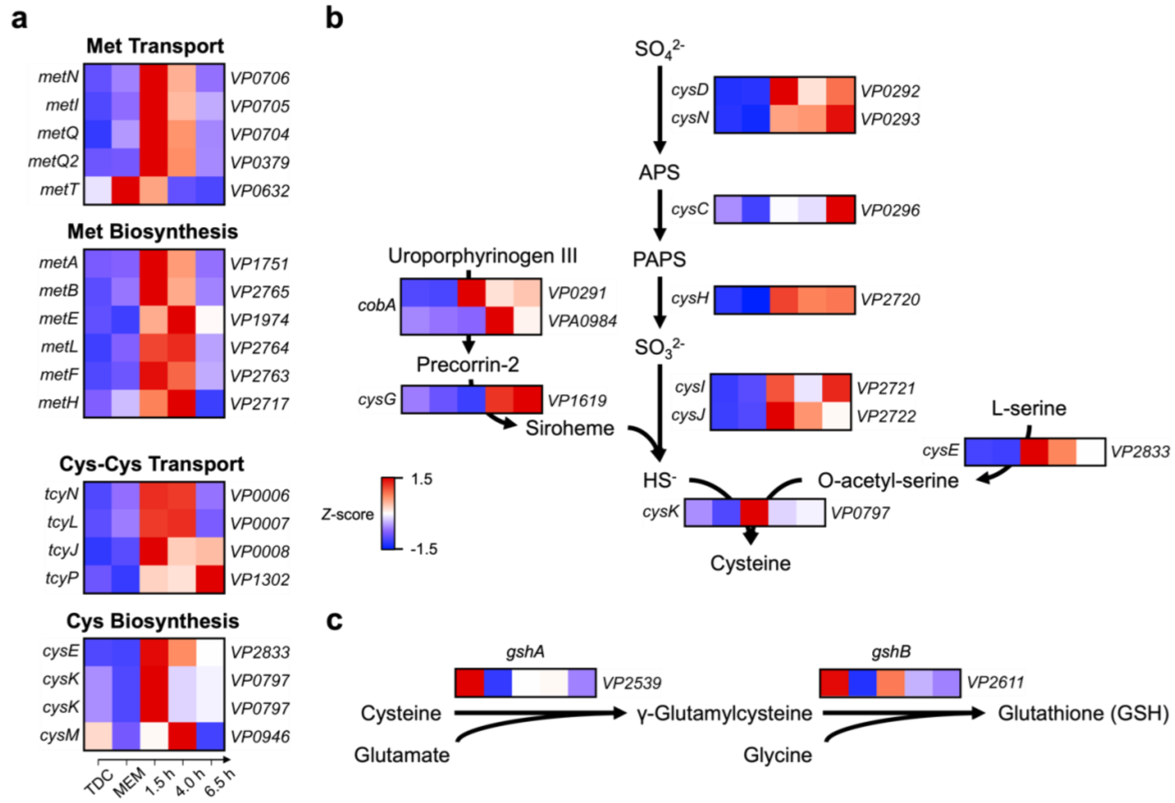


**Figure S5. Early induction of sulfur metabolism in *V. para* during intracellular infection.**

**(a)** *Z*-score-normalized expression of genes involved in the biosynthesis and transport of sulfur-containing amino acids, including methionine (Met), cysteine (Cys), and cystine (Cys-Cys), across pre-infection and intracellular infection conditions. Heatmaps show mean expression across two biological replicates, with gene names on the left and locus tags on the right.

**(b)** *Z*-score-normalized expression of genes involved in sulfate assimilation across pre-infection and intracellular infection conditions. Heatmaps are presented as in **a**. APS, adenosine 5′-phosphosulfate; OAS, O-acetylserine; PAPS, 3′-phosphoadenosine 5′-phosphosulfate.

**(c)** *Z*-score-normalized expression of genes involved in glutathione biosynthesis across pre-infection and intracellular infection conditions. Heatmaps are presented as in **a**.

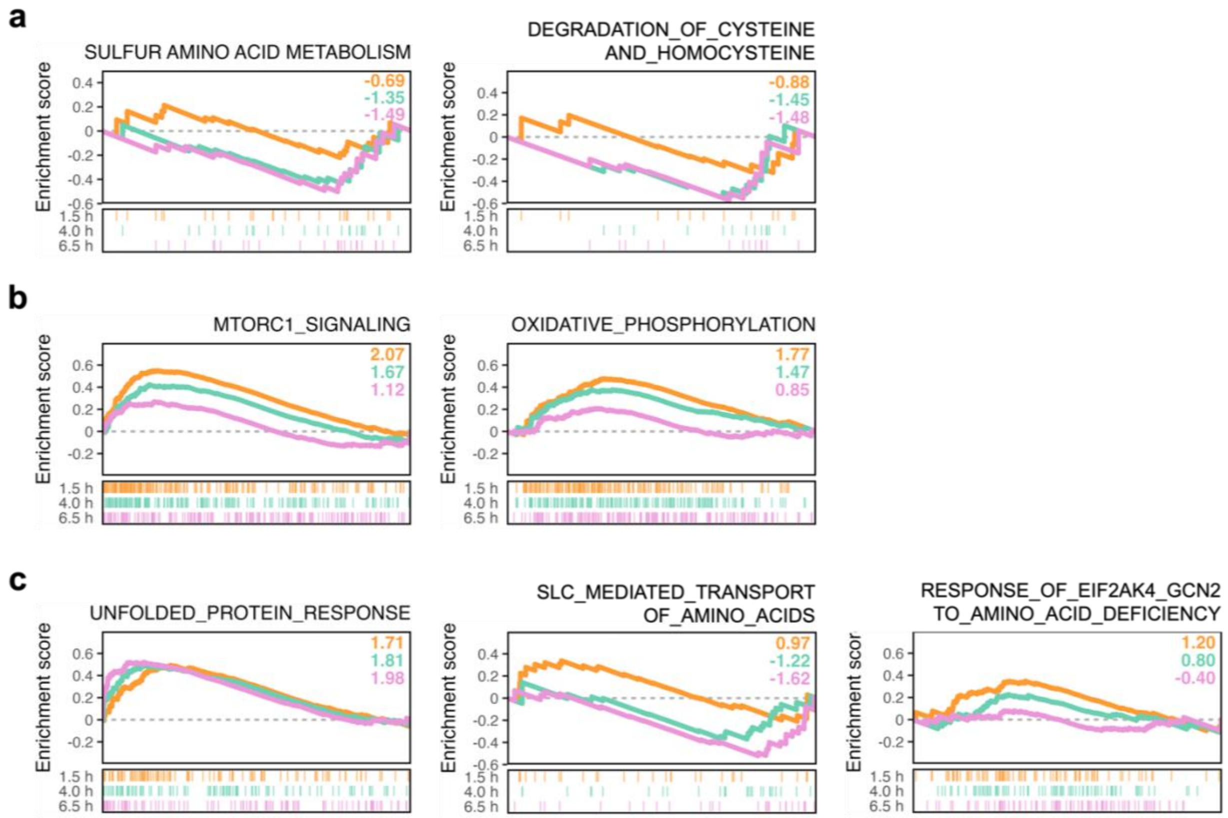


**Figure S6. Temporal remodeling of host metabolic pathways during *V. para* infection.**

GSEA running-score plots of pathways associated with **(a)** sulfur amino acid metabolism, **(b)** metabolic signaling and energy metabolism, and **(c)** amino acid transport and cellular stress responses. All pathways were analyzed in infected host cells relative to uninfected mock controls. Normalized enrichment scores (NES) are indicated in each plot. Colors denote infection time points: orange, 1.5 h; teal, 4.0 h; and pink, 6.5 h, shown from top to bottom. MTORC1_SIGNALING, OXIDATIVE_PHOSPHORYLATION, and UNFOLDED_PROTEIN_RESPONSE were obtained from the Hallmark gene-set collection; all other pathways were obtained from Reactome.

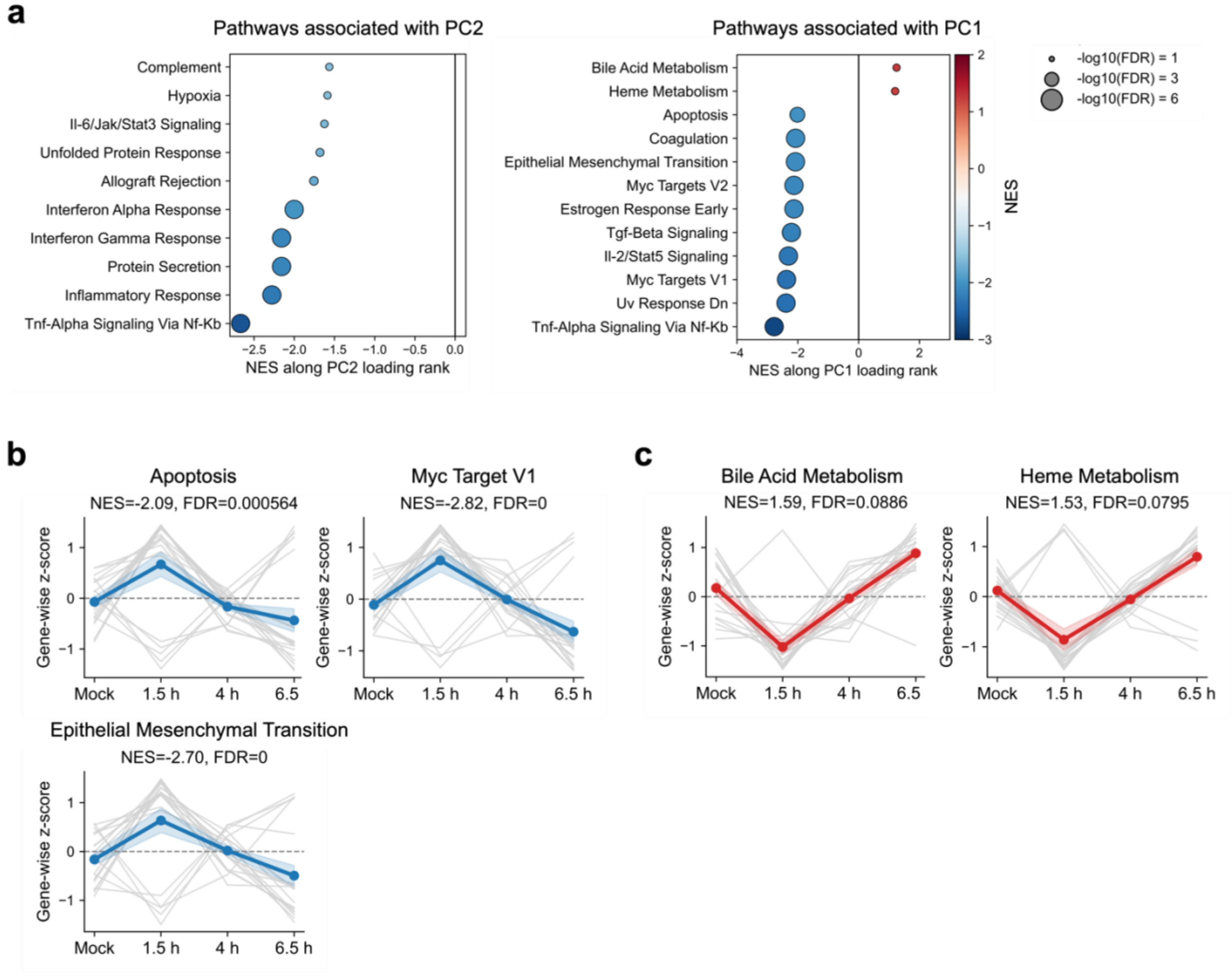


**Figure S7 Host pathways associated with PC1 and their temporal expression trajectories.**

**(a)** Hallmark pathways associated with the major principal components. GSEA was performed using genes ranked by their loadings along each principal component (PC). PC1 and PC2 are shown on right and left panel, respectively. The top positively and negatively enriched Hallmark gene sets with FDR $q < 0.25$ are shown. Each dot represents one gene set. The x-axis indicates the normalized enrichment score (NES), with positive and negative values indicating enrichment toward the positive and negative ends of the PC gene-loading rank, respectively. Dot size represents −log10(FDR $q$-value), and dot color indicates NES.

**(b–c)** Temporal expression trajectories of the top 20 genes from selected PC1-associated pathways. For each gene, expression values were log2-transformed, averaged across biological replicates within each condition, and standardized as $z$-scores across mock, 1.5, 4, and 6.5 h. Gray lines show the trajectories of individual genes, while the colored line represents the mean trajectory across the 20 genes; shading indicates the SEM across genes. Blue and red denote pathways with negative and positive NES values, respectively. NES and FDR $q$-values are indicated above each plot.

### Supplementary Table Captions

**Table S1. Pathway enrichment analysis of *V. para* temporal gene-expression clusters.**

Functional enrichment results for the seven gene-expression clusters identified during intracellular infection, as shown in **Fig. 2c**. Annotations include KEGG pathways, GO Biological Process terms, RegPrecise-predicted regulons, and manually curated gene sets. Enrichment was assessed using hypergeometric tests. Benjamini–Hochberg-adjusted P values are reported for GO terms, whereas nominal P values are reported for other gene sets.

**Table S2. Curated annotation of *V. para* nutrient-uptake systems and their predicted substrate classes.**

Nutrient transport systems identified in the *V. para* genome and presented in **Fig. 3c**. Genes predicted to transport the same substrate are grouped in the same row and annotated according to their substrate categories.

**Table S3. Mean TPM values of *V. para* secretion system components across infection conditions.**

Mean transcript abundance (TPM) of genes encoding secretion system components, effectors, and chaperones under pre-invasion (TDC and MEM) and intracellular infection (1.5 h, 4 h, and 6.5 h) conditions. Values represent the means of two biological replicates.

**Table S4. Primers used for construction of *V. para* strains and complementation plasmids.**

Primer sequences used for generating bacterial deletion mutants and their corresponding complementation constructs.

**Table S5. Customized oligonucleotide probes used for *V. para* rRNA depletion.**

Sequences of custom oligonucleotide probes designed to target *V. para* ribosomal RNA during dual RNA-seq library preparation.

**Table S6. Primers and oligonucleotides used for dual RNA-seq library construction.**

Sequences of primers and oligonucleotides used for reverse transcription, amplification, and sequencing library preparation.
